# Aberrant activation of wound healing programs within the metastatic niche facilitates lung colonization by osteosarcoma cells

**DOI:** 10.1101/2024.01.10.575008

**Authors:** James B. Reinecke, Leyre Jimenez Garcia, Amy C. Gross, Maren Cam, Matthew V. Cannon, Matthew J. Gust, Jeffrey P. Sheridan, Berkley E. Gryder, Ruben Dries, Ryan D. Roberts

## Abstract

**Purpose:** Lung metastasis is responsible for nearly all deaths caused by osteosarcoma, the most common pediatric bone tumor. How malignant bone cells coerce the lung microenvironment to support metastatic growth is unclear. The purpose of this study is to identify metastasis-specific therapeutic vulnerabilities by delineating the cellular and molecular mechanisms underlying osteosarcoma lung metastatic niche formation.

**Experimental design:** Using single-cell transcriptomics (scRNA-seq), we characterized genome- and tissue-wide molecular changes induced within lung tissues by disseminated osteosarcoma cells in both immunocompetent murine models of metastasis and patient samples. We confirmed transcriptomic findings at the protein level and determined spatial relationships with multi-parameter immunofluorescence and spatial transcriptomics. Based on these findings, we evaluated the ability of nintedanib, a kinase inhibitor used to treat patients with pulmonary fibrosis, to impair metastasis progression in both immunocompetent murine osteosarcoma and immunodeficient human xenograft models. Single-nucleus and spatial transcriptomics was used to perform molecular pharmacodynamic studies that define the effects of nintedanib on tumor and non-tumor cells within the metastatic microenvironment.

**Results:** Osteosarcoma cells induced acute alveolar epithelial injury upon lung dissemination. scRNA-seq demonstrated that the surrounding lung stroma adopts a chronic, non-resolving wound-healing phenotype similar to that seen in other models of lung injury. Accordingly, metastasis-associated lung demonstrated marked fibrosis, likely due to the accumulation of pathogenic, pro-fibrotic, partially differentiated epithelial intermediates and macrophages. Our data demonstrated that nintedanib prevented metastatic progression in multiple murine and human xenograft models by inhibiting osteosarcoma-induced fibrosis.

**Conclusions:** Fibrosis represents a targetable vulnerability to block the progression of osteosarcoma lung metastasis. Our data support a model wherein interactions between osteosarcoma cells and epithelial cells create a pro-metastatic niche by inducing tumor deposition of extracellular matrix proteins such as fibronectin that is disrupted by the anti-fibrotic TKI nintedanib. Our data shed light on the non-cell autonomous effects of TKIs on metastasis and provide a roadmap for using single-cell and spatial transcriptomics to define the mechanism of action of TKI on metastases in animal models.

**Statement of translational relevance:** Therapies that block metastasis have the potential to save the majority of lives lost due to solid tumors. Disseminated tumor cells must integrate into the foreign, inhospitable microenvironments they encounter within secondary organs to facilitate metastatic colonization and progression. Our study elucidated that disseminated osteosarcoma cells survive within the lung by co-opting and amplifying the lung’s endogenous wound-healing response program. This osteosarcoma-induced wound response results in fibrosis of the surrounding microenvironment. Our data implicates fibrosis and abnormal wound healing as key drivers of osteosarcoma lung metastasis that can be targeted therapeutically to disrupt metastasis progression.

## Introduction

Metastasis is responsible for nearly all solid tumor-related deaths [1]. Metastasis proceeds through a multistep process wherein tumor cells invade the surrounding normal tissue adjacent to the primary tumor, enter the blood stream (for tumors that spread via hematogenous route), colonize, and ultimately grow within a distant tissue [2]. Preclinical and patient-level data suggest that therapies designed to disrupt the early steps of the metastatic cascade, such as invasion through adjacent tissues and intravasation into the bloodstream, are less likely to prevent clinically-relevant metastatic lesions, as disseminated malignant cells have likely colonized distant tissues prior to the detection of a primary tumor [3–5]. Development of therapies that target the later elements of the metastatic cascade are more likely to have clinical impact but have remained largely theoretical due to our crude understanding of the mechanisms that drive the progression of tumor growth in distant tissues following colonization. Understanding how disseminated tumor cells transform the foreign and otherwise hostile microenvironments of metastatic sites into accommodating environments that promote tumor growth may expose novel metastatic vulnerabilities [6].

The lung is a common site of metastasis for tumors that originate in diverse developmental origins [7]. Physiologically, the lung not only facilitates gas exchange, but also creates a barrier to the invasion of microbes, toxins, pollutants, and allergens. Accordingly, the lung has evolved complex mechanisms for recognizing and repairing injury without causing excessive tissue damage or inflammation [8]. Certain pathological conditions such as idiopathic pulmonary fibrosis (IPF) arise when the mechanisms of lung repair become altered. In these altered states, dysregulation of these wound healing mechanisms leads to progressive dysfunction [9]. At the cellular level, IPF is characterized by epithelial hyperplasia, accumulation of fibroblasts, and the excessive deposition of extracellular matrix (ECM) proteins such as fibronectin and collagen, all confined within an immunosuppressive microenvironment [10]. How these structural elements of the lung respond to metastatic cancer cells remains poorly understood. The profoundly inefficient nature of the metastatic process and the dramatic differences evident in the comparison of normal lung to metastatic lesions suggest that the soil of the lung must be shaped by cancer cells in order for tumors to successfully colonize and grow in the organ [11]. Therefore, disrupting the tumor education of the lung microenvironment is an attractive therapeutic approach to targeting metastatic disease.

Osteosarcoma is the most common malignant bone tumor in young people and ostensibly metastasizes to the lung, leading to poor patient outcomes [12–14]. Given the organotropism that osteosarcoma displays for the lung, and the critical clinical problem lung metastasis poses to patients with osteosarcoma, we leveraged several murine and human xenograft models to elucidate how the lung responds to metastatic osteosarcoma cells. We uncovered that osteosarcoma cells arrest within the distal alveolar epithelial niche, damage alveolar epithelial cells, and trigger an abnormal alveolar wound healing program that promotes metastatic progression. Utilizing single-cell RNA sequencing (scRNA-seq) and spatial transcriptomic analysis, we found that the stroma of established metastases resembles a fibrotic, non-healing wound. Translationally, disrupting the lung fibrosis response with nintedanib, a tyrosine kinase inhibitor (TKI) utilized clinically for treating patients with IPF, slowed osteosarcoma metastasis by reprogramming the metastatic microenvironment. Collectively, our study sheds light on how tumors co-opt the lung microenvironment during metastasis and provides proof-of-concept that targeting the metastatic niche is a viable therapeutic strategy.

## Materials and methods

### Cell culture

All cell lines utilized herein were routinely authenticated with STR and mycoplasma testing through Laboratory Corporation of America. Cells utilized for experiments were within 20 passage doublings after reviving from liquid nitrogen storage. OS-17 cells were derived from an early passage of the OS-17 patient-derived xenograft created from a primary femur biopsy performed at St. Jude’s Children’s Research Hospital in Memphis and was a gift from Peter Houghton. K7M2 cells are a highly metastatic derivative cell line derived from a spontaneous osteosarcoma in a BALB/c mouse and were obtained from the American Type Culture Collection (ATCC, CRL2836). F420 cells are a murine osteosarcoma derived from spontaneous lung metastases arising in the genetically engineered osteosarcoma mouse model Col2a-Cre mutant Tp53 and was a gift from Jason Yustine. HBEC3-KT (HBEC) is an immortalized non-tumorigenic human lung epithelial cell line and were obtained from the ATCC (CRL-4051). F420 and K7M2 cells were propagated in culture in DMEM (Corning, 10-013-CV) supplemented with 10% fetal bovine serum (FBS) (R&D Systems, S11150H) while OS-17 cells were cultured in RPMI (Corning, 10-040-CV) supplemented with 10% FBS. HBEC cells were propagated in human airway epithelial media (ATCC, PCS-300-030) with the Bronchial Epithelial Cell Growth Kit (ATCC, PCS-300-040). OS-17 and HBEC cells were plated 1:1 on to glass coverslips in DMEM/F12 (Corning, 10-092-CV) supplemented with 1% FBS. HBEC conditioned media was obtained after 72 hours of incubation (DMEM/F12 with 1% FBS) and was cleared with centrifugation at 1000 rpm prior to treating osteosarcoma cells.

### Animal studies

All animal studies carried out in this study were done with the approval of the Institutional Animal Care and Use Committee of the Research Institute at Nationwide Children’s Hospital (IACUC protocol AR15-00022) prior to conducting experiments. Six- to eight-week-old female C57BL/6 (for F420; RRID: MGI:2159769), BALB/c (for K7M2; RRID: IMSR_APB:4790), and CB17-SCID (for human OS-17 xenograft- CB17SC scid-/- (RRID: IMSR_TAC:cb17sc)) mice were used for all studies. For generation of primary tumors, single cell suspensions of 5×10^5^ cells were injected intra-tibially. Primary tumors were excised once they grew to 800 mm^3^. For generation of experimental metastasis, F420 (5×10^5^), K7M2 (5×10^5^), or OS-17 (1×10^6^) cells were inoculated intravenously via tail vein. Cell number utilized for experimental metastasis is titrated by model to ensure survival of mice post injection and to ensure metastasis rates of at least 80%[15]. Mice were followed until designated endpoint of experiment or until arriving at humane endpoint. For nintedanib (MedChemExpress, HY-50904) treatment, experimental metastasis was induced as described above. To determine the effect of nintedanib on metastatic progression, one day after injection, mice were randomized to receive vehicle (0.5% hydroxyethylcellulose, Eisen-Golden Laboratories, EG-HEC-3) or nintedanib 50 mg/kg twice per day. Mice were monitored and once an ill mouse (which was a vehicle treated mouse in all three models) was confirmed to have lung metastasis, all vehicle and nintedanib treatment mice were sacrificed. All mice were euthanized by IACUC approved CO_2_ inhalation method prior to processing as described below. To define the effects of nintedanib on established metastases using single-nucleus transcriptomics, mice were injected with F420 cells expressing luciferase. The development of metastatic disease was monitored weekly by intra-peritoneal injection of luciferin followed by bioluminescence imaging. On day 28 post injection, asymptomatic mice with identifiable metastasis were randomized to receive vehicle or nintedanib 50 mg/kg BID X14 days and processed for single-nucleus RNA sequencing or histology as described below.

### Tissue dissociation, single-cell and single-nucleus RNA library preparation, and sequencing

Lungs harvested from mice were processed using the human tumor dissociation kit (Miltenyi Biotec, 130-095-929) with a GentleMacs Octo Dissociator with Heaters (Miltenyi Biotec, 130-096-427) for 3’ single-cell analysis. For single-nucleus, cryopreserved metastasis-bearing lungs were isolated and permeabilized for single-cell gene expression sequencing following the Nuclei Isolation from Complex Tissues for Single Cell Multiome ATAC + Gene Expression Sequencing protocol (10X Genomics, CG000375 Rev C). Following dissociation into single cells or nuclei, 3’ RNA-Seq libraries were prepared using Chromium Single Cell 3′RNA-sequencing system (10X Genomics) with the Reagent Kit v3.1 (10X Genomics, PN-1000121) according to the manufacturer’s instructions. cDNA library quality and quantity were determined using an Agilent 2100 bioanalyzer using the High Sensitivity DNA Reagents kit (Agilent, 5607-4626) and then sequenced on an Illumina NovaSeq 6000. Raw data and initial processing was performed using cellranger version 7 (10X Genomics) with reads mapped to mm10 and hg19 versions of the mouse and human genomes.

### Visium spatial library preparation and sequencing

A histological section was generated from a mouse with lung metastases from injected F420 tumor cells as outlined above. This section was stained with hematoxylin and eosin and imaged using NIS-Elements AR software (v. 5.42) using a Nikon Eclipse Ti2-E microscope with a Nikon DS-Ri2 color camera and a 10x Plan Apochromat Lambda D objective. The section was then mounted into a CytAssist instrument (10X Genomics) and spatial libraries were prepared according to the manufacturer’s recommendations. The resulting library was sequenced on an Illumina NextSeq 2000. The SpaceRanger software (10X Genomics, version 2.1.1) was used to align the data to the mm10 mouse genome, assigning reads to both genes and spots on the Visium slide and aligning the data to H&E images.

### Single-cell and single-nucleus RNA analysis

All code used for data analyses is available on GitHub (https://github.com/kidcancerlab/epithelial).

10X single-cell and single-nuclei sequencing data were loaded into Seurat toolkit version V5 in R for further analysis [16]. Cells were filtered by minimum and maximum UMI counts appropriate to each sample (available in the GitHub repository) and any cell with mitochondrial DNA >20% was removed. RNA counts were normalized with regularized negative binomial regression via SCTransform. After dimensionality reduction with principal component analysis and UMAP embedding, unsupervised clustering was achieved using “FindNeighbors” and “FindClusters” function in Seurat using PCA dimensions 1:30. Clustering resolution was determined using the clustree R package[17]). Differential gene expression between clusters was determined using FindMarkers (Seurat) and the Wilcox method. For single-cell data, cell types were approximated with SingleR using the Human Lung Cell Atlas and the Human Primary Cell Atlas as references [18, 19]. For single-nuclei data, we used publicly available single-nuclei reference data as a reference[20]. Clusters for both types of data were then curated manually based on the automated assignments and review of canonical marker expression.

Human normal lung single-cell data were obtained from the Human Lung Cell Atlas, with samples filtered using the accompanying metadata to isolate cells from healthy lungs, nonsmokers, lung parenchyma, and then downsampled in a weighted fashion to include similar proportions of whole cell and nuclear libraries as the lung metastasis samples [18].

Differentially expressed genes were evaluated by GSEA against KEGG, GO:Molecular Function, GO:Biological Processes, and transcription factor activity (all from migdb) using fgsea [21]. Results were plotted in R using the ggplot suite.

The nichenetr R package was used to identify cell-cell interactions within the tumor microenvironment in our single-cell RNAseq data that might be disrupted by nintedanib treatment[22]. To identify all potential interactions, each cell type was, in turn, considered a “receiver” cell type while the other cell types were “sender” cells producing ligands. For each receiver cell type, gene expression fold change values were calculated for genes up-regulated in tumor-bearing lung compared to healthy lung.

Potential ligands and receptors for each cell-cell interaction were selected by removing those not expressed in sender and receiver cells, respectively. The output from nichenetr was filtered for receptors potentially inhibited by nintedanib to generate a list of ligand-receptor interactions to assess in samples treated by nintedanib[23, 24]. The nintedanib-treated samples were single-nuclei and so were sub-optimal to perform nichenetr analyses in directly due to a lack of steady-state cytoplasmic RNAs. However, we were able use these samples to assess the downstream targets of the ligand-receptor interactions identified in single-cell data as outlined above. Briefly, the databases provided with the nichenetr package were used to identify all genes predicted to be downstream of the ligand-receptor interactions identified in the single-cell data as potentially inhibited by nintedanib. These gene lists were then used as input for the AUCell R package to generate activation scores for each nuclei in the dataset[25]. The distribution of these activation scores were plotted as violin plots for the untreated F420-bearing lung as well as the F420-bearing lungs treated with nintedanib at either 3- or 28-days post tumor tail vein injection. To assess differences between groups we calculated FDR-adjusted *p*-values using t-tests and Cohen’s d statistics as an estimate of effect size.

### Visium spatial data analysis

Visium spatial RNA sequencing data was loaded into Seurat V5 in R for further analysis. Data outside of the tissue was filtered and the remaining data was normalized using SCtransform and then principal component analysis, cluster assignment and further dimensional reduction by UMAP was performed. The single-cell data generated by this study was used as a reference to deconvolute each spot in the spatial dataset using the spacexr R package[26].

To assess activity of pathways potentially inhibited by nintedanib (as identified in single-cell data as outlined above) we used the AUCell R package to generate activation scores for each spot in the Visium data. These activation scores were then plotted to show spatial patterns of potential pathway activation in tumor-bearing lung.

### Histology, immunohistochemistry, and tissue clearing

Sternotomy was performed and lungs were flushed via slow instillation of phosphate buffered saline (PBS) (Corning, 21-031-CV) through right ventricle until lungs were cleared of blood. Lungs were dissected from mouse *en bloc* along with trachea and heart. Lungs were slowly instilled with 10% neutral buffered formalin (NBF) (Sigma-Aldrich, HT5012-1CS) until fully expanded, then trachea was tied off with suture and placed in NBF for 24 hours at 4°C. After repeated washing in PBS, lung lobes were carefully dissected from each other and processed for paraffin embedding (FFPE) or immunohistochemistry frozen (IHC-F). For FFPE, 4µm sections were deparaffinized with xylene and rehydrated through ethanol series. For cryosections, lung lobes were cryoprotected through a sucrose gradient (10%, 15%, 30%), mounted in Optimum Cutting Temperature compound (Fisher Scientific, 23-730-571), frozen in liquid nitrogen and then sectioned to 10µm. To quantify metastasis burden, FFPE sections were counterstained with hematoxylin (Sigma-Aldrich, MHS16-500ML) and eosin (Abcam, AB246824) via conventional methods. Heat-mediated antigen retrieval methods utilized citrate buffer (CB; pH 6.9) or Tris-EDTA buffer (TE; pH 9) with Tween 20 (Fisher Bioreagents, BP337-100) at 0.2% v/v. Antigen retrieval (AR) buffer used was antibody dependent (see below).

For thick tissue analysis, Ce3D-mediated tissue clearing was performed as described previously[27, 28].

### Immunofluorescence and microscopy

FFPE sections were rehydrated in PBS for 30 minutes. Human tissue sections utilized in this study were obtained from Nationwide Children’s Pathology department under institutional review board approved study (IRB1100478). Cells growing on #1.5 glass coverslips (Electron Microscopy Sciences, 72204-04) were fixed in NBF for 10 minutes at room temperature. Following three rinses in PBS, coverslip or histological sections were then incubated for at least 1 hour in blocking solution consisting of PBS, 0.2% triton100 (v/v) (Sigma, X-100-100ml), and 3% bovine serum albumin (w/v) (Sigma, A-7888) at room temperature. For Ce3D specimens, blocking and permeabilization was performed overnight at room temperature. Primary antibodies used: rabbit anti-fibronectin (Abcam, ab2413, AR=TE), rabbit anti-mouse Sftpc (Abcam, ab211326, AR= CB or TE), rabbit anti-mouse Ly6G (CST, 87048, AR=CB), hamster anti-Podoplanin (Developmental Studies Hybridoma Bank(DSHB), 8.1.1, AR=TE), rat anti-KRT8 (DSHB, TROMA-I, AR=TE), rat anti-Ki67 (Invitrogen, 14-5698-82, AR=CB), rabbit anti-human SFTPC (Millipore, ABC99, AR=TE), mouse anti-human fibronectin (R&D,MAB1918, AR=TE), rabbit anti-mouse Cd68 (R&D, MAB101141, AR= CB or TE). Primary antibodies were diluted in blocking solution and sections were incubated in primary antibody solution overnight at 4°C. For Ce3D, primary antibody incubation was performed for 72 hours 4°C. Following three rinses in PBS (for Ce3D, 3 washes x 2 hours/wash), sections were incubated with appropriate AlexaFluor labeled secondary antibodies (Invitrogen), DAPI (Invitrogen, D1306), and where indicated, Phalloidin AlexaFluor 647 (Invitrogen, A12379) or Wheat Germ Agglutinin (WGA) AlexaFluor 647 (Invitrogen, W11261) diluted in blocking solution for 1 hour at room temperature. For Ce3D, tissue was incubated for 24 hours at room temperature. Following PBS rinses, coverslips and tissue sections were mounted in Fluoromount G (Invitrogen, 00495802).

### Microscopy and image analysis

Fluorescent microscopy images were obtained on a LSM 800 confocal microscope using 20X air or 63X oil objectives. Tile-scanning was used for high-resolution imaging of large areas with images stitched together with 10% overlap using ZenBlue software. CZI files were then uploaded and processed using ImageJ. All manipulation of images (brightness/contrast and smoothing) were done uniformly to each image. For quantifying Krt8 and fibronectin fluorescence, samples were processed and imaged simultaneously under same microscope settings, then identical manual threshold was applied to all images in ImageJ. Fluorescence was measured/area with “measure” analysis function. Krt8 expression was normalized to Krt8 fluorescence in areas without visible metastases.. The cell counter plugin was used to manually quantify Ki67+/ Sftpc+ cells, Ly6G+ neutrophils, or CD68+ macrophages. Metastasis burden was quantified from H/E sections by blinded examiner using Aperio ImageScope Software (V12.4.3). Whole lung lobe images were obtained with Aperio FL ScanScope digital slide scanner with 20X objective.

### Statistical analysis

Statistical analysis for animal and bench studies was performed using Graphpad Prism V9. The number of samples and mice are explicitly stated in text with appropriate statistical test and *p*-values. Data reported with error bars represent mean +/- standard error of the mean (SEM). Statistical analyses of the sequencing data are described in the relevant sections.

### Data availability and resource sharing

Code used to analyze single-cell RNA-seq (scRNA-seq) data can be found on GitHub at github.com/kidcancerlab/epithelial. Previously unpublished raw sequencing data and the associated count tables with metadata is available through GEO accession GSE252703. Human data is available on dbGaP though accession phs003569.v1.p1 and GEO accession GSE270231. Cell lines utilized in this study are available from commercially available resources or can be obtained upon request to corresponding author.

## Results

### Disseminated osteosarcoma cells localize to alveolar epithelial niche and disrupt alveolar architecture during metastatic progression

We first sought to determine the spatial localization of osteosarcoma cells as they begin to colonize the lung, which would provide evidence for the specific lung environment most relevant to these earliest steps in metastatic colonization. To control the timing of dissemination, we inoculated immunocompetent C57BL/6 mice intravenously (tail vein) with F420 murine osteosarcoma cells. We then collected lungs from mice euthanized at 1, 7, and 14 days after injection and processed lungs onto slides to evaluate the development of the earliest metastatic niche. F420 cells demonstrated a stereotyped pattern of lung colonization characterized by extravasation and invasion through the alveolar epithelial layer, inducing injury to podoplanin (PDPN)+ type 1 alveolar epithelial cells (AEC1). We identified a compensatory increase of surfactant protein C (Spc)+ type 2 alveolar epithelial cells (AEC2) over areas of injured AEC1 (Figure 1A), reminiscent of the process triggered by other types of alveolar injury [29] . Similar results were seen in the K7M2 murine osteosarcoma model (Supplemental Figure 1A).

**Figure 1:**
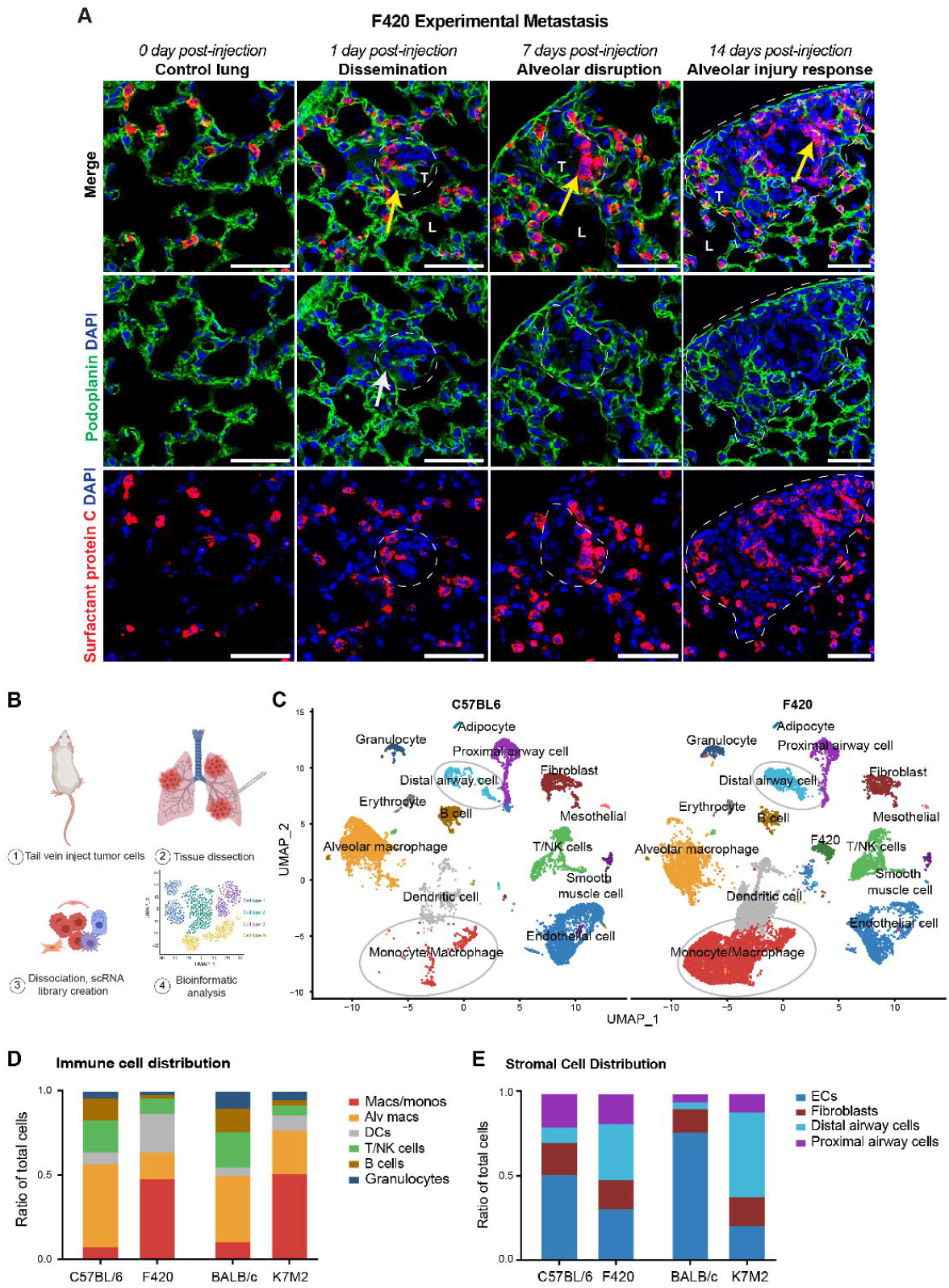
Osteosarcoma lung colonization induces acute alveolar injury response. **A**, IHC of control lung and early metastatic lesions (marked by dash circle). White arrow denotes AEC1 (Podoplanin;green) injury while yellow arrow notes AEC2 response (Spc;red). Nuclei are counterstained with DAPI (blue). Scale bar= 50µm. **B**, Schematic for approach to metastasis scRNA-sequencing. **C**, Uniform manifold projection (UMAP) demonstrating cell clusters in non-metastasis bearing C57BL6 compared to F420 metastasis. Grey circles denote populations enriched in metastasis bearing lung **D-E**, Relative quantification of cell type distribution by scRNA-sequencing.

In other nonmalignant models of alveolar injury, disruption of alveolar integrity promotes neutrophil followed by macrophage infiltration into the lung[30]. We likewise observed a short burst of neutrophil infiltration shortly after colonization that diminished over time and was followed by a significant and sustained accumulation of macrophages (Supplemental Figure 1B), mirroring the kinetics of immune infiltration observed in other alveolar injury models [30]. Taken together, these data suggested that disseminated osteosarcoma cells colonize the distal alveolar niche and induce an acute lung injury response during early metastatic colonization.

To understand how these changes affect the behavior of the surviving osteosarcoma cells and the surrounding lung tissues, we isolated early F420 and K7M2 metastases using a stereomicroscope and processed them for single-cell RNA sequencing (scRNA-seq; Figure 1B). Cell cluster annotation was done using a combination of automated and manual approaches using established markers (Supplemental Figure 2A;C, methods). Multiple cell populations demonstrated enrichment in metastasis when compared to non-tumor bearing control lung tissues (Figure 1C, Supplemental Figure 2B). We divided cells computationally into infiltrating/immune and native/stromal compartments to quantify the relative frequencies of each distinct cell population. We stress that this quantification has inherent bias in the capture of certain cell populations by droplet based scRNA-seq methods [31]. With this caveat in mind, we found that metastasis bearing lungs exhibit a marked enrichment in infiltrating myeloid cells (particularly macrophages) and distal airway epithelial cells (Figure 1D-E). Immunofluorescence imaging confirmed that developing metastases become highly enriched in macrophages and distal airway epithelial cells when compared to uninvolved lung (Supplemental Figure 2D-E). We further confirmed the enrichment of epithelial cells in metastasis compared to unaffected lung areas using the thick-tissue Ce3D tissue clearing method (Supplemental Video 1-2). We next isolated the alveolar epithelial cells and macrophages to evaluate the phenotypic changes in the two most clearly altered stromal and immune cell populations, hypothesizing that the invasion of the osteosarcoma cells influences these changes, and that these constitute important elements of tumor education within the developing metastatic niche.

### Metastasis associated epithelial cells adopt a non-healing wound phenotype

Distal epithelial cell clusters from metastases and healthy (non-metastasis bearing) C57BL/6 (control for F420) or BALB/c (control for K7M2) mice were merged, integrated, and re-analyzed through our Seurat-based pipeline. Several populations of epithelial cells were enriched in metastasis bearing lungs compared to control (Figure 2A). Five distal epithelial clusters were identified in F420 and K7M2 metastases, each with distinct gene expression signatures (Figure 2B).

**Figure 2:**
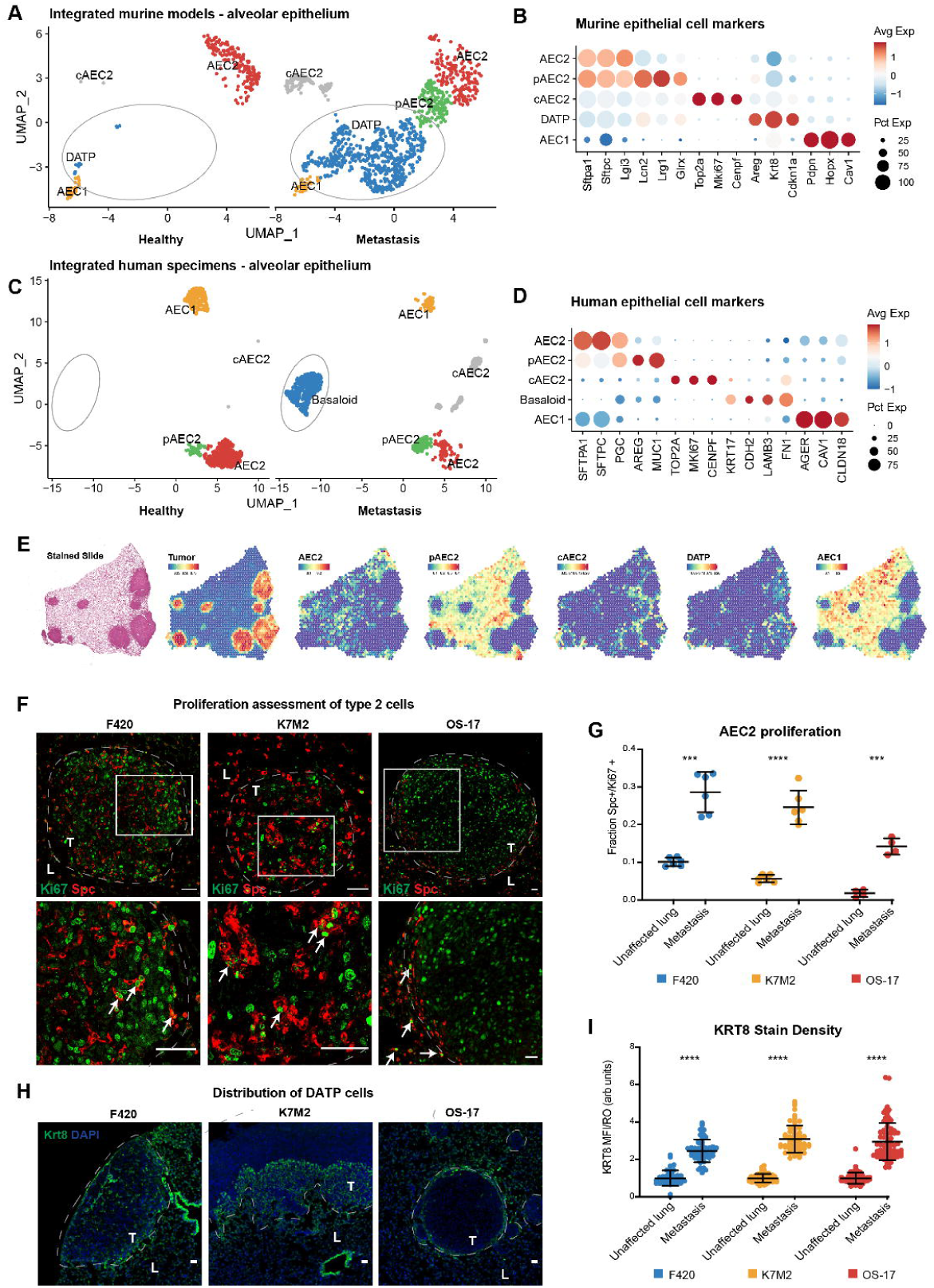
Metastasis associated epithelial cells acquire chronic wound phenotype. **A**, UMAP plot of murine alveolar epithelial cells in healthy lung compared to metastasis. Grey circle denotes marked enrichment of DATP in metastasis. **B**, Dotplot demonstrating expression of key marker genes (columns) in alveolar epithelial cell clusters (row). **C**, UMAP plot of normal human lung specimens compared to patient lung metastasis samples. Grey circle denotes enrichment of basaloid cells in osteosarcoma metastasis. **D**, Dotplot demonstrating expression of key marker genes. **E**, Visium spatial transcriptomic analysis illustrating localization of alveolar epithelial cell subsets enriched in a metastasis-bearing murine lung. **F**, IHC of Spc (red; AEC2 marker) and Ki67 (green; proliferation marker). L= unaffected lung, T= metastasis. Dash circle notes lung-metastasis border. Rectangle marks inset shown below. Areas demonstrate examples of Spc+Ki67+ cells. Nuclei are counterstained with DAPI (blue). Scale bar= 50µm. **G**, Quantification of AEC2 proliferation. 100 cells/animal, n=4. Welch’s t-test: F420 p= 0.0003, K7M2 p= <0.0001, OS-17 p= 0.0004. **H**, IHC of Krt8 (green; DATP marker). Scale bar= 50µm. Dash circle notes lung-metastasis border. Areas demonstrate examples of Krt8+ cells around growing metastatic lesions. Nuclei are counterstained with DAPI (blue). Scale bar= 50µm. **I**, Quantification of DATP accumulation (by Krt8+ fluorescence). N=4 animals, at least 10 regions of interest per animal for metastasis and unaffected lung. Welch’s t-test: p=<0.0001.

AEC1 and AEC2 populations were discernible based on the enrichment of genes such as *Cav1*, *Hopx* and *Sftpc* respectively. Two additional AEC2-like populations were found within the metastatic samples, one of which expressed genes such as *Lcn2* and *Lrg1* while the other demonstrated upregulation of genes involved in cell cycle such as *Mki67* and *Cdk1*. This finding was consistent with the physiologic reaction to alveolar injury, as AEC2 cells are facultative stem cells within the alveolus, functioning to re-establish epithelial barrier integrity after AEC1 injury [29]. In the canonical wound healing process, AEC2 are activated into a ‘primed’ pAEC2 state by AEC1 injury [32]. pAEC2 then enter the cell cycle (cAEC2) before differentiating into AEC1 cells to re-establish alveolar epithelial integrity. Given their phenotypic resemblance of these previously-defined subtypes, we assigned the *Lcn2*^+^ population a label of ‘primed’ AEC2 (pAEC2) and the *Mki67*^+^ cluster a label of ‘cycling’ AEC2 (cAEC2), consistent with the established literature for murine lung injury models [32].

The fifth epithelial cluster demonstrated upregulation of atypical cytokeratins, such as *Krt8* and *Krt18/19*, and genes such as *Cdkn1a*, *Sfn*, and *Nupr1.* This was an interesting finding, as Krt8^+^ is a primary marker of a recently described fibrogenic intermediate produced as activated AEC2 cells differentiate into AEC1 cells in response to injury [32–34]. Under normal conditions, these intermediates are short-lived and do not persist beyond the acute phase of the injury. However, under certain chronic inflammatory conditions, these damage associated transient progenitors (DATP) (also referred to as alveolar differentiation intermediates (ADI) and pre-alveolar type-1 transitional cell state (PATS)) become arrested in the Krt8+ transitional state and can promote pathologic fibrosis. Indeed, a similar process has been shown to occur in human patients with chronic idiopathic pulmonary fibrosis [32–34].

To ensure that these findings reflect biology that is relevant to the human disease, we collected tissue samples from patients undergoing metastatectomy at our local institution and performed single-cell sequencing using the same protocol. To enhance the power of this analysis and ensure the broad applicability of these findings, we combined these with previously published datasets containing both normal lung tissues and metastatic osteosarcoma lesions [18, 19, 35–37]. This assessment of human tissues revealed changes in both epithelial content and phenotype, very similar to those observed in our murine models, with several epithelial cell clusters enriched in metastasis relative to healthy lung (Figure 2C-D).

Of note, in human lung pathologic conditions, AEC2 have been shown to differentiate through a disease-specific, aberrant KRT17+ basaloid cell state. These KRT17+ cells stain positive for KRT8 by immunohistochemistry (IHC) and cluster with murine Krt8+ DATP in cross-species single-cell transcriptomic analyses [34, 38]. Thus, they are likely to be an orthologous AEC intermediate. Our human osteosarcoma metastasis datasets replicated the finding in the murine models, with strong enrichment of KRT17+ basaloid cells (Figure 2C-D).

We next performed spatial transcriptomics to localize epithelial subsets relative to metastatic lesions in the F420 immunocompetent murine model (Figure 2E). Epithelial subsets like cAEC2 and DATP were only found in proximity to metastatic lesions, supporting that they are metastasis-induced cell types. pAEC2 and cAEC2 localized in the metastasis:lung border while DATP cells infiltrated into metastatic lesions. We confirmed these sequencing and spatial transcriptomic findings in tissue slices by showing that AEC2 within the developing metastatic niches were hyperproliferative (Ki67+) relative to AEC2 in uninvolved areas of the lung across multiple models of osteosarcoma metastasis (Figure 2F-G). Additionally, accumulation of metastasis-associated Krt8+ epithelial cells occurs across multiple osteosarcoma models (Figure 2H-I). Lastly, KRT8+ AEC intermediates were prevalent within human lung metastasis samples by immunofluorescence (Supplemental Figure 3). These findings demonstrate that basaloid/DATP accumulation is a conserved phenomenon in osteosarcoma metastasis and not unique to mouse models.

We conclude that partially differentiated epithelial intermediates, which have been associated with aberrant non-healing lung wounds in other pathological processes, consistently accumulate within the developing metastatic niche in both mice and humans. Functionally, this finding suggested that impaired epithelial wound healing is central to osteosarcoma lung metastasis. As lung alveolar wound repair is a highly stereotyped and regulated process that requires coordination with other cell types (such as fibroblasts and macrophages [39]), we wondered whether the monocyte-derived macrophages that accumulate around developing metastatic lesions might evidence changes characteristic of either physiologic or pathologic wound healing. We thus leveraged our single-cell datasets to define the macrophage phenotypes associated with osteosarcoma metastasis.

### Metastasis associated macrophages are enriched in scar, chronic wound-like phenotype that are spatially segregated in osteosarcoma metastasis

Macrophages can play paradoxical roles during distinct phases of wound healing [40]. In the early wound healing response, macrophages are predominantly inflammatory in nature [41, 42], marked by high expression of antigen presentation proteins and inflammatory cytokines in the early phases of wound healing [43]. Once the inciting noxious stimulus is controlled, anti-inflammatory/resolution macrophages promote healing by reorganizing the extracellular matrix, phagocytosing apoptotic cells, and dampening inflammation, though their uncontrolled accumulation can promote chronic, non-healing wounds [44].

To ascertain the macrophage phenotypes associated with developing osteosarcoma metastases, we isolated myeloid cells from our murine and human scRNA-seq datasets. In this analysis, we also included primary murine tibial tumors to define how the surrounding microenvironment impacts macrophage phenotypes in the primary vs. metastatic microenvironment. Murine osteosarcoma metastases demonstrated marked accumulation of several macrophage populations that were transcriptionally distinct from populations present in healthy lung and tibial tumors (Figure 3A-B). Tumor associated macrophage annotations were made using recently published criteria [45]. Notably, a population of macrophages expressing *Trem2*, *Cd9*, *and Spp1* were highly enriched in lung metastasis compared to healthy lung and primary tibial tumors. This macrophage population shares transcriptional profile with scar-like macrophages[46], a macrophage phenotype characterized by expression of genes such as *Cd9, Trem2, Spp1, Gpnmb,* and *Fabp*5, which have recently been identified as a crucial driver of pathologic fibrosis within the lung and during liver metastasis [47–49]. Their enrichment in osteosarcoma lung metastasis, but not in primary tibial tumors, suggests a microenvironmental influence on macrophage phenotypes, with chronic injury/fibrosis being promoted specifically in the lung, likely secondary to osteosarcoma-induced aberrant wound healing process. Similar macrophage subpopulations were enriched in human osteosarcoma metastasis relative to healthy human lung (Figure 3C-D).

**Figure 3:**
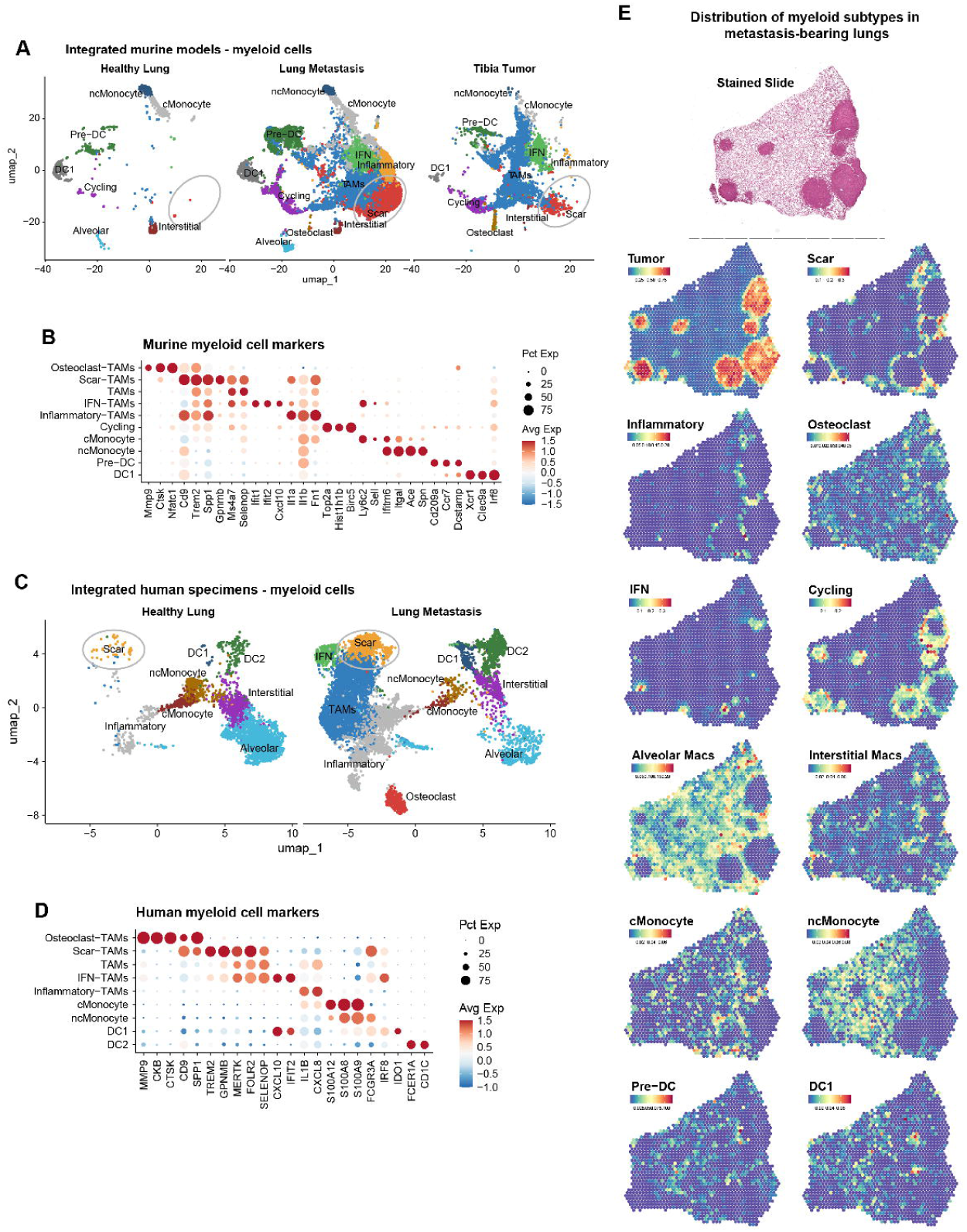
Metastasis associated macrophages adopt wound-like phenotypes that are spatially segregated. **A**, UMAP of myeloid cells including macrophages and dendritic cells in healthy lungs compared to F420 metastasis and F420 tibia tumor. Grey circle notes enrichment of scar macrophages in lung metastasis relative to tibia tumor and healthy lung. **B**, Dotplot demonstrating expression of key marker genes (columns) in myeloid cell clusters (row). **C**, UMAP plot of normal human lung specimens compared to patient lung metastasis samples. Grey circle denotes enrichment of scar macrophages in osteosarcoma metastasis. **D**, Dotplot demonstrating expression of key marker genes. **E**, Visium spatial transcriptomic analysis illustrating localization of myeloid cell subsets and demonstrating the spatial enrichment of specific macrophage subsets at the tumor-lung border in a metastasis-bearing murine lung.

Given the complexity of macrophage phenotypes and overlapping marker expression, we utilized spatial transcriptomics to define the localization of fibrotic macrophage subsets relative to established metastases (Figure 3E). Scar macrophages were only localized to the metastatic compartment of the lung, like epithelial fibrotic cell types, and confined to the lung:metastasis border. Spatial analysis also demonstrated marked enrichment of actively proliferating (cycling) macrophages, inflammatory macrophages, as well as recruitment of classical monocytes to metastatic lesions relative to unaffected lung. Based on our epithelial and macrophage data, we propose a model wherein the activation of epithelial wound healing, triggered by the infiltrating osteosarcoma cells at the initial stages of metastasis and continuing along the leading edge during lesional growth, coincides with the enrichment of inflammatory macrophages. Subsequently, the tumor-educated inability to resolve the wound causes macrophages with chronic wound phenotypes to accumulate within the metastatic lesion. In other disease states, this chronic wound healing process ultimately leads to pulmonary fibrosis, characterized by the pathological deposition of extracellular matrix proteins such as fibronectin [50]. Therefore, we next investigated if osteosarcoma metastases are fibrotic.

### Osteosarcoma cells co-opt epithelial:mesenchymal interactions to promote fibrotic reprogramming of metastasis associated lung

Fibronectin is an extracellular matrix protein that is necessary for proper wound healing but can promote progressive organ dysfunction if not tightly regulated [51]. In all three of our osteosarcoma models tested, fibronectin markedly accumulated within developing metastatic lesions, relative to adjacent unaffected lung. Given the findings above, this is consistent with a metastasis-induced fibrotic transformation of the osteosarcoma associated lung (Figure 4A). To elucidate whether osteosarcoma-associated fibronectin deposition is metastasis-specific, we compared primary F420 and K7M2 tibial tumors to lung metastases. Primary tibial tumors expressed low levels of intracellular fibronectin while metastases were characterized by extensive extracellular, fibrillar fibronectin matrices at the lung-metastasis interface, suggesting that fibronectin matrix formation is metastasis specific and not an inherent trait of osteosarcoma tumors (Figure 4B).

**Figure 4:**
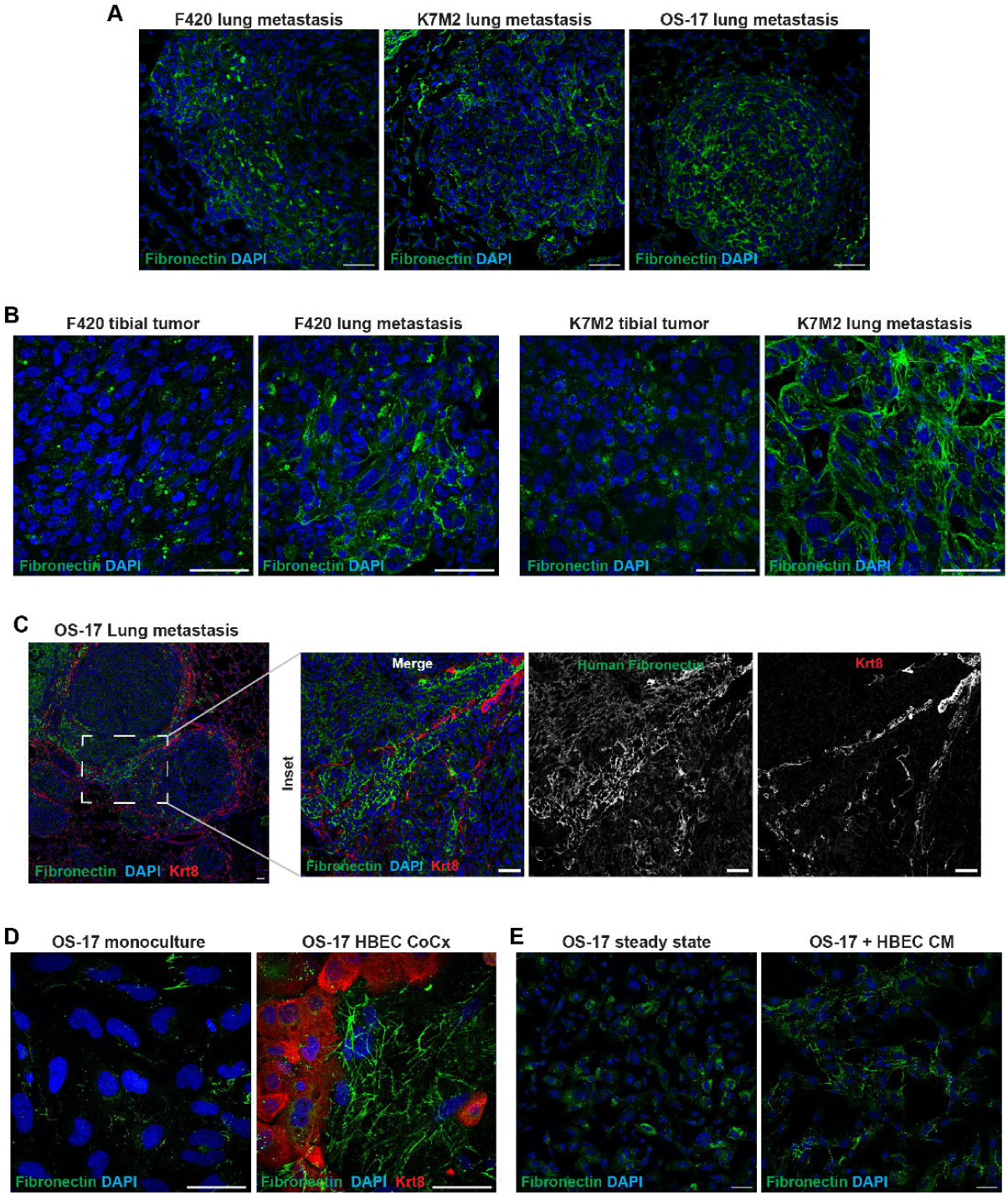
Lung epithelial cells induce metastasis-specific osteosarcoma fibronectin matrix formation. **A,** IHC for fibronectin in several osteosarcoma metastasis models. **B,** IHC comparing fibronectin staining in tibial tumors vs lung metastasis. **C,** IHC for fibronectin (green) and Krt8 (red; DATP cells). Dotted box notes inset, which demonstrates area of metastasis enriched in Krt8+ DATP cells and human (osteosarcoma) fibrillar fibronectin. **D,** Immunofluorescence of fibronectin matrix in OS-17 cells cultured alone or in coculture with human bronchial epithelial cells (HBEC; KRT8=Red). **E,** Immunofluorescence of fibronectin in OS-17 cells cultured at steady state or stimulated with conditioned HBEC media. Scale bar= 50µm. Nuclei are counterstained with DAPI (blue).

In non-malignant lung fibrosis diseases, activated myofibroblast secrete and organize fibronectin into dense matrices [52]. Additionally, fibronectin deposition by cancer-associated fibroblasts is also noted in other tumor types [53]. However, our scRNA-seq data did not demonstrate enrichment of myofibroblasts in osteosarcoma metastases, suggesting an alternative source of fibrotic matrix in osteosarcoma metastases. Given the mesenchymal origin of osteosarcoma, it is developmentally related to fibroblasts and demonstrates strong matrix-forming tendencies. We leveraged the xenogenic nature of the OS-17 metastasis model to help determine whether metastasis-associated fibronectin is tumor-cell derived. To do this, we stained metastatic lesions with a human-specific fibronectin antibody. Human (tumor) derived fibronectin was highly expressed throughout metastatic lesions. However, extracellular matrix fibrils were enriched at the lung-metastasis interface, suggesting that local microenvironment interactions favor extracellular matrix deposition at this location.

In lung fibrotic disorders, epithelial damage triggers fibronectin secretion and fibrosis by stimulating mesenchymal cells [54]. Based on our data demonstrating osteosarcoma cells induce epithelial injury, we hypothesized that epithelial cells may induce osteosarcoma fibronectin deposition. Indeed, in our OS17 model, fibronectin matrix deposition correlated with dense collections of Krt8+ DATP cells (Figure 4C). To test this hypothesis within a reductionist system, OS-17 cells were co-cultured with human immortalized lung epithelial cells (HBEC) or stimulated with epithelial conditioned media. OS-17 cells generated a complex fibronectin matrix in the presence of lung epithelial cells (Figure 4D). This reaction was replicated by stimulation with epithelial-conditioned media (Figure 4E), suggesting that the reaction is triggered by soluble signals produced by the epithelial cells.

In summary, chronic unresolved injury induced by osteosarcoma lung colonization culminated in microenvironmental fibrosis. This fibronectin-rich matrix was synthesized by osteosarcoma cells in a metastasis-dependent manner due to microenvironment specific epithelial interactions. We thus hypothesized that osteosarcoma-induced chronic injury and subsequent fibrotic transformation of the lung niche are key drivers of osteosarcoma metastasis progression and that inhibiting this metastasis-associated fibrosis would prevent osteosarcoma metastasis growth.

### Nintedanib, an anti-fibrotic tyrosine kinase inhibitor, inhibits metastasis formation

Fibrotic lung conditions are progressive, often fatal diseases. Nintedanib is an orally available TKI with a broad target profile that has demonstrated clinical efficacy in slowing the progression of fibrotic lung diseases [24, 55, 56]. The pathophysiological overlap with osteosarcoma metastases and lung fibrotic conditions suggested that anti-fibrotic agents such as nintedanib could disrupt metastasis progression. To test the effect of nintedanib on osteosarcoma metastasis progression, tumor cells were injected intravenously via tail vein to promote experimental metastasis formation. Nintedanib or vehicle was initiated at 24h after injection to focus on the impact of nintedanib on metastasis progression, not extravasation into lung (Figure 5A). Study mice were monitored daily for signs of metastasis. When metastasis was confirmed in one mouse, all mice were humanely euthanized and lung metastases quantified (Figure 5B-G). We found that nintedanib significantly hindered metastatic progression in both immunocompetent (F420 and K7M2) and immunodeficient (OS-17) models.

**Figure 5:**
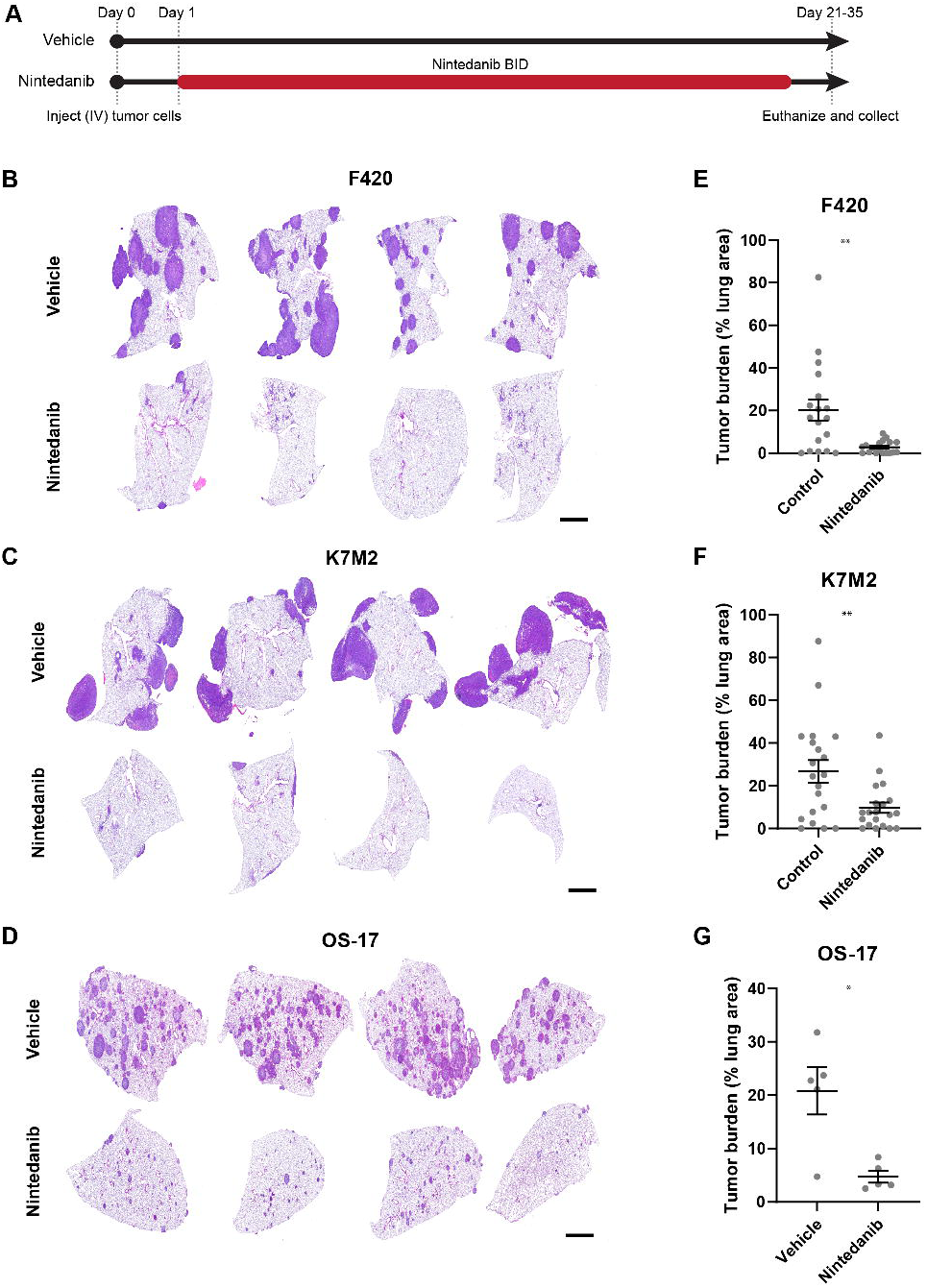
Anti-fibrotic tyrosine kinase inhibitor Nintedanib disrupts metastatic progression. **A,** Schematic of nintedanib trial. **B-D**, Hematoxylin and eosin staining of representative images from 4 animals for each treatment group. Scale bar= 1000µm. **E-G**, Quantification of tumor burden/lung lobe in vehicle vs nintedanib treated mice. F420 and K7M2 = 20/treatment group, OS-17= 5/treatment group. Welch’s t-test F420 p= 0.0030, K7M2 p= 0.0068, OS-17= 0.0199.

Interestingly, nintedanib did not significantly affect F420 or OS17 cell viability *in vitro* (Supplemental Figure 4A). Additionally, nintedanib did not affect primary tibial tumor growth in a F420 model while causing slowing but not regression of OS17 tibial growth (Supplemental Figure 4B-D). Collectively, the conserved activity of nintedanib in disrupting metastasis progression across models and within three distinct mouse strains strongly supports the anti-metastatic effect in osteosarcoma. Given nintedanib’s clinical activity in fibrotic diseases, and its marked effect on metastatic osteosarcoma compared to primary tibial tumors or cells growing *in vitro*, we surmised that nintedanib likely inhibited metastatic progression by disrupting the osteosarcoma-induced, pathologically aberrant wound-healing process within the lung.

### Nintedanib prevents osteosarcoma-induced chronic wound phenotype

Nintedanib is a promiscuous TKI with documented activity on multiple receptor tyrosine kinase receptors such as FGFR1-4, VEGFR, and PDGFRα/β [23, 24]. We next utilized a combination of spatial and single-cell transcriptomics to gain insights into the mechanisms underlying the potent effects of nintedanib on osteosarcoma lung metastasis. We devised an experimental approach using NicheNet, a bioinformatic tool that identifies transcriptomic changes induced by ligand-receptor interactions, to identify transcriptional activation of nintedanib target pathways in the metastatic lung relative to unaffected lung. We then performed snRNA-seq on mice treated with nintedanib to identify cell types and pathways impacted by nintedanib during metastasis progression (immediate treatment) or on established metastases (delayed treatment; Figure 6A). First, we projected nintedanib target pathways identified by NicheNet analysis (Supplemental Figure 5A-) onto our spatial data set to identify where nintedanib targets are active in metastatic lesions. Strikingly, while activation of pathways such as FGFR and PDGFR was markedly upregulated in metastatic lesions relative to unaffected lung, FGFR activation was spatially restricted to the lung:metastasis border while PDGFR was preferentially activated in the metastasis core (Figure 6B).

**Figure 6:**
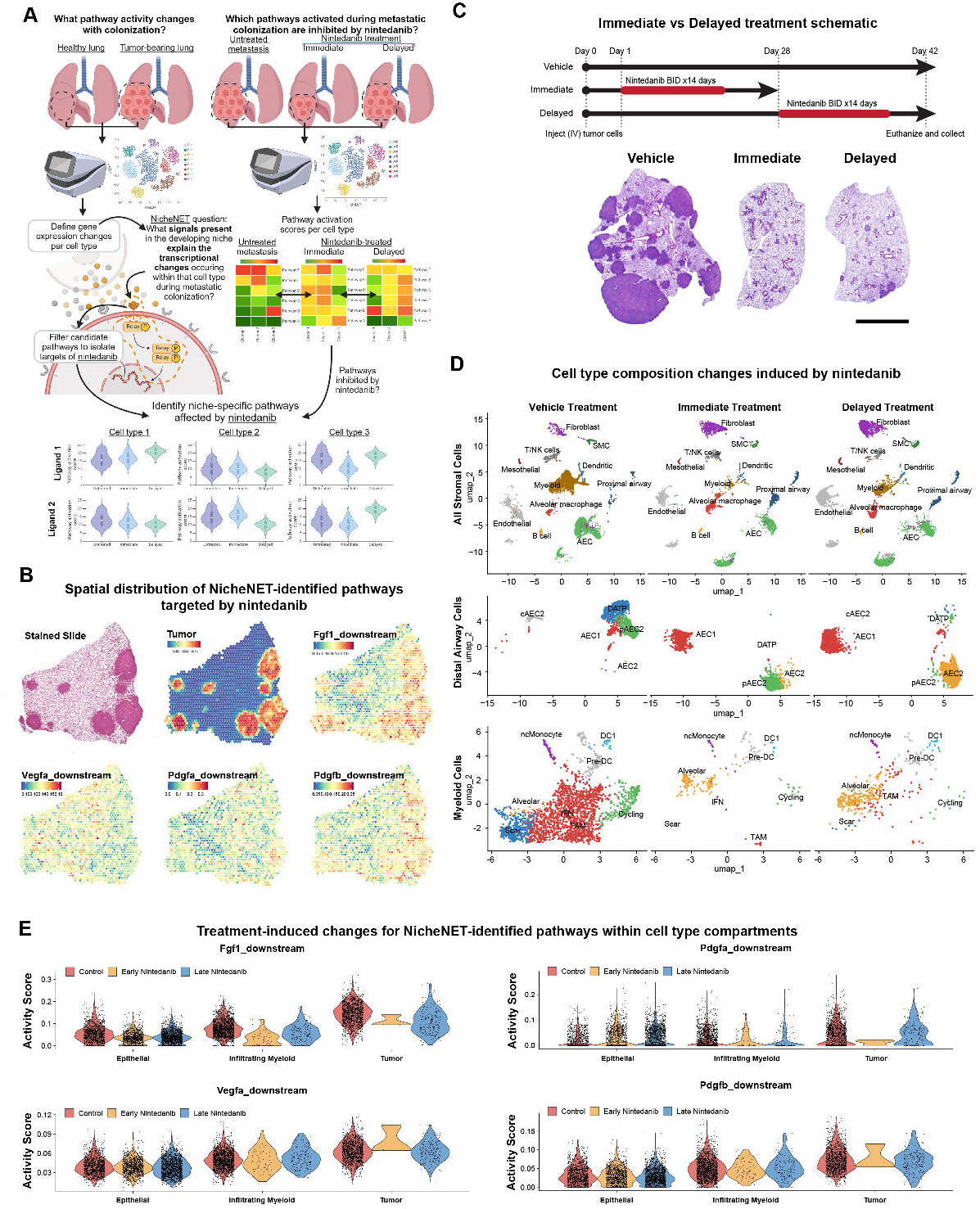
Nintedanib blocks essential signals in both tumor and niche cell compartments. **A,** Schematic of bioinformatic analysis of single cell gene expression data to identify niche-specific pathways affected by nintedanib. **B,** Visium spatial transcriptomic analysis demonstrating expression of identified nintedanib-targeted signaling pathways in a tumor-bearing murine lung. **C,** Schematic of nintedanib regimen. Hematoxylin and eosin staining of representative images from an animal for each treatment group. Scale bar= 1000µm. **D,** UMAP plots of cell composition in untreated, immediate, and delayed nintedanib-treated F420 metastases. A subset of cell clusters is highlighted to denote the changes in two cell populations of interest: Distal airway cells and macrophages. **E,** Violin plots of activity scores for nintedanib-targeted niche-specific pathways identified within cell type compartments of interest.

To understand how nintedanib affects the cell types that constitute the metastatic niche and the pathways impacted by nintedanib, we treated mice with nintedanib shortly after metastatic colonization (1 day after injection) or once metastases were established and confirmed by bioluminescence imaging (28 days after injection, Figure 6C demonstrates treatment schematic and representative H/E images for each treatment group; see Supplemental Figure 6 for bioluminescence imaging). Single-nucleus transcriptomic analysis demonstrated that nintedanib prevented the accumulation of fibrotic DATP and scar-macs in the immediate treatment group, consistent with nintedanib inhibiting metastasis-induced fibrosis (Figure 6D). Remarkably, nintedanib also reversed metastasis-induced fibrotic phenotypes, as evidenced by a decrease in DATP and scar-macs in the delayed treatment group (Figure 6D). Lastly, we validated the effect of nintedanib at the protein level *in vitro* and *in vivo*, demonstrating that nintedanib inhibited epithelial-induced tumor fibronectin deposition as well as decreasing fibronectin and DATP cells in established metastases *in vivo* (Supplemental Figure 6A-D). We concluded that nintedanib fundamentally alters the cellular and matrix composition of osteosarcoma metastases, impacting both tumor cells and surrounding stromal cells.

We next utilized our NicheNet results to ask what nintedanib targeted pathways are downregulated by nintedanib in key cell types: tumor cells, infiltrating myeloid cells, and epithelial cells (Figure 6E). FGFR activation was downregulated by nintedanib in all three cell types, while the impact on PDGFR was limited to tumor cells and infiltrating myeloid cells. VEGFR signaling was not impacted in these three cell types, which is consistent with VEGFR signaling being primarily restricted to endothelial cells (Supplemental Figure 8). Collectively, the results of our preclinical nintedanib trial, coupled with this molecularly-guided pharmacodynamic analysis of nintedanib-treated metastases, suggest a tight linkage between the anti-metastatic activity and the anti-fibrogenic activity of nintedanib. These results also suggest a requirement for fibrotic reprogramming of the lung microenvironment during osteosarcoma metastasis progression. Inhibition of this fibrogenic, aberrant wound healing program represents a novel therapeutic vulnerability with the potential for combatting osteosarcoma lung metastasis.

## Discussion

How disseminated cancer cells colonize, integrate with, and coerce foreign organs to promote metastatic growth remains a poorly understood phenomenon. With respect to osteosarcoma, specifically, our data supports a model wherein the tumor cells trigger and then perpetuate the lung’s intrinsic wound healing program, beginning with dissemination and resulting in a chronic, non-healing fibrotic wound. The characterization of a tumor as a non-healing wound is not new. It was first postulated by Virchow, then further conceptualized by Dvorak. However, this paradigm largely focused on tumorigenesis of primary tumors in adult epithelial carcinomas [57]. Our results demonstrate that osteosarcoma, a mesenchymal pediatric tumor, induces alveolar epithelial injury upon colonization of the lung, leading to a chronic non-healing wound and fibrotic transformation of the surrounding lung. In many ways, osteosarcoma-epithelial interactions mimic, albeit in a highly pathological sense, epithelial-lung fibroblast interactions that occur during non-malignant lung wounds [54, 58]. This observation may suggest that aberrant epithelial-mesenchymal interactions are central to lung metastasis across diverse histological subtypes.

Indeed, an analogous scenario has been demonstrated in breast cancer lung metastasis, wherein epithelial tumor cells co-opt and activate normal lung mesenchymal fibroblasts to drive metastasis, with fibrogenic wound-like processes playing a central role in the mechanisms for colonization [59]. It may be that in the case of breast cancer metastasis, the tumor cells play the role of the epithelial cells, and the fibroblasts the role of the mesenchymal cell in a process similar to that observed here. If true, this aberrant wound-healing process and the resulting fibrogenic changes may represent mechanisms broadly conserved across tumor types, though this requires rigorous and direct exploration across tumor models. Notably, organs with facultative repair capacity that are susceptible to fibrosis, such as the lung, liver, and pancreas, are organs commonly affected by high metastatic rates (or, in the case of the pancreas, characterized by fibrotic/desmoplastic primary tumors) [8, 60, 61]. It will be critical from a therapeutic perspective to elucidate whether initiation of abnormal wound healing and progression to organ fibrosis is specific to a handful of histological tumor types or a generalizable mechanism.

Our work significantly advances the mechanistic understanding of osteosarcoma lung metastasis. Our findings converge on previous findings by Zhang and colleagues but add several critical new observations [62]. Notably, our work demonstrates that fibrogenic reprogramming is a niche-wide event initiated by osteosarcoma-epithelial interactions and provides critical pharmacodynamic analysis of nintedanib-treated metastases in an immunocompetent host. Our mechanisms also unify and advance concepts emerging from other recent publications, suggesting that cancer cells disrupt the physiology of the lung epithelia during colonization [63, 64]. Long considered inert bystanders in the processes that facilitate malignant progression, stromal cells such as alveolar epithelial cells have received relatively little attention as potential drivers of metastasis biology. However, our work establishes the key role that epithelial cells play in metastatic colonization. Indeed, the clear physical proximity and interactions of early disseminated osteosarcoma cells with alveolar epithelial cells, together with the functional data demonstrating that epithelial cells initiate fibrogenic reprogramming of osteosarcoma cells, strongly suggest active two-way paracrine signaling between osteosarcoma cells and epithelial cells as a key event driving osteosarcoma colonization of the lung. Future work is required to clarify the mediators of these osteosarcoma-epithelial interactions.

The use of TKIs in sarcoma patients is well-established, especially in the relapse setting [65, 66]. Indeed, many have been evaluated, several of which have partially overlapping targets with nintedanib. For instance, cabozantinib is an orally available TKI that inhibits VEGFR2, c-MET, and RET and has activity in the relapsed/metastatic setting [67]. Nearly 40-50% of osteosarcoma demonstrate amplification of VEGFA [68], and MET signaling can transform osteoblasts [69], the likely cell of origin for osteosarcoma. Accordingly, cabozantinib could work on both the microenvironment (inhibiting angiogenesis) and tumor cells (inhibiting MET signaling), but likely would not inhibit tumor or stromal FGFR and PDGFR signaling as implicated by our single-cell and spatial transcriptomic analysis. To our knowledge, the tumor cell-intrinsic and -extrinsic mechanisms mediating the anti-metastasis effects of cabozantinib and other clinically-validated TKIs remain poorly understood. Our data suggest that nintedanib exerts its effect in a context-dependent manner (lung metastasis) and provide evidence that nintedanib’s anti-metastatic activity likely involves suppression of fibrogenic signaling pathways involving both tumor and host cells, particularly those involving FGFR-induced transcriptional responses. Even in the treatment of pulmonary fibrosis, the most clinically important target(s) of nintedanib remain unclear. Nintedanib’s spectrum of activity includes inhibition of VEGFR, PDGFR, and FGFR signaling, as well as several intracellular kinases such as SRC, which is critical for the signal transduction of many RTKs inhibited by nintedanib [23]. This target promiscuity is common amongst the TKIs, including those already used to treat osteosarcoma [70]. While several of these targets are shared, this work provides an initial suggestion that the unique targets may account for the clinical effects of individual agents and that the on-target, off-tumor effects may be just as important as those intrinsic to the tumor cells. Here, we demonstrated how single-cell and spatial transcriptomics can be leveraged to further our mechanistic understanding of the holistic impact that therapeutic interventions have on the metastatic microenvironment. Given that TKIs often demonstrate activity against unanticipated targets [71], a more comprehensive evaluation using such approaches may point to novel mechanisms and suggest strategic combinations of agents that would otherwise not be pursued.

Elucidating why fibronectin/fibrosis is required for osteosarcoma metastatic progression remains an important area of ongoing investigation. Potential, non-mutually exclusive mechanisms include: directly providing survival signals to tumor cells during early lung colonization [72], resistance to chemotherapy [73, 74], protection against innate/adaptive anti-tumor immunity [75], and reprogramming of niche cells such as macrophages. We note that the effectiveness of nintedanib was similar in mice with or without intact adaptive immunity, which does not rule out an important role for fibrosis in controlling anti-tumor immunity but does support the idea that fibronectin/fibrosis could function through both tumor cell-intrinsic and non-tumor cell-intrinsic mechanisms to promote metastatic progression.

In conclusion, we note that metastases are complex multi-cellular structures composed of tumor, immune, and altered resident stromal cells. Utilizing scRNA-seq and spatial transcriptomics, we illuminate how osteosarcoma cells manipulate the lung microenvironment into a chronic state of wound healing. From a translational perspective, we demonstrate that targeting the microenvironment is a viable therapeutic option across multiple osteosarcoma models, which is an important consideration due to the substantial intertumoral heterogeneity inherent to osteosarcoma.

## Acknowledgements

We would like to thank the patients and their families for donation of tissue. We would like to thank the Histopathology, Microscopy, Biopathology Center Processing and Banking, and Institute for Genomic Medicine sequencing cores at Nationwide Children’s Hospital for their technical assistance. This work is supported by Alex’s Lemonade Stand Crazy 8 Award (RDR,RD,BEG), Alex’s Lemonade Stand Foundation Young Investigator Award (JBR), NIH R01CA260178 (RDR), NIH CTSA Grant UL1TR002733 (RDR), Translational Studies Grant from the Osteosarcoma Institute (RDR), CancerFree KIDS Foundation (RDR, JBR), Steps For Sarcoma Foundation (RDR,JBR), Sarcoma Foundation of America (RDR), Nationwide Children’s Director’s Strategic Development Fund (RDR).

