## Supplemental figures 1-8 for "Aberrant activation of wound healing programs within the metastatic niche facilitates lung colonization by osteosarcoma cells"

Supplemental Figure 1: K7M2 shows evidence of acute alveolar injury during osteosarcoma metastasis

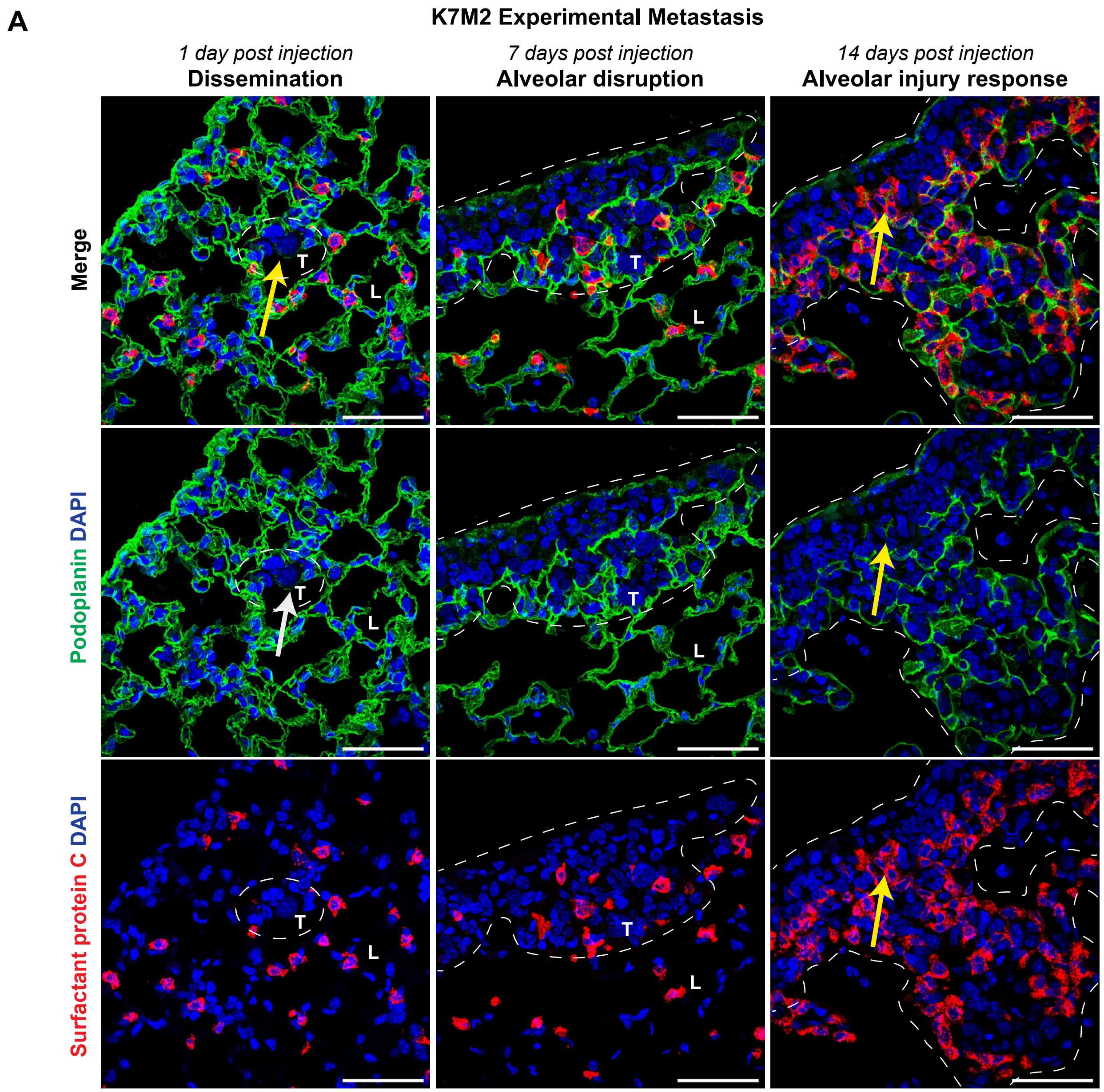


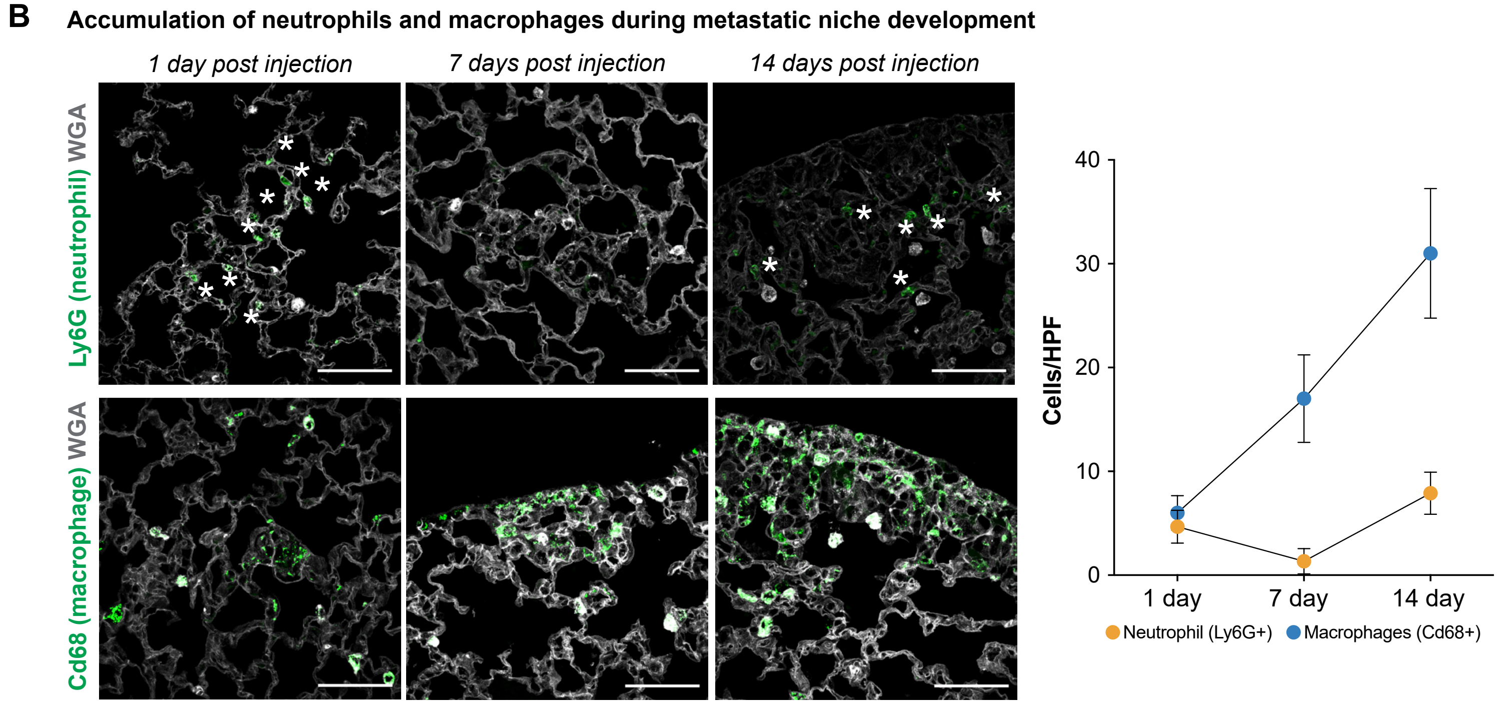

**Supplemental Figure 1:** Alveolar injury response in K7M2 murine osteosarcoma model. **A,** IHC of control lung and early metastatic lesions (marked by dash circle). White arrow denotes AEC1 (Podoplanin;green) injury while yellow arrow notes AEC2 response (Spc;red). Scale bar= 50μm. **B,** IHC demonstrating kinetics of neutrophil (Ly6G+) and macrophage (Cd68+) recruitment during early metastatic progression. Wheat Germ Agglutinin (WGA;grey) marks lung parenchyma. Asterisks denote Ly6G+ neutrophils. Nuclei are counterstained with DAPI (blue). Scale bar= 50μm. Neutrophil and macrophage quantitation, n=3 animals per timepoint.

**Supplemental Figure 2: scRNA-seq marker expression and protein level validation of macrophage and epithelial cell enrichment**

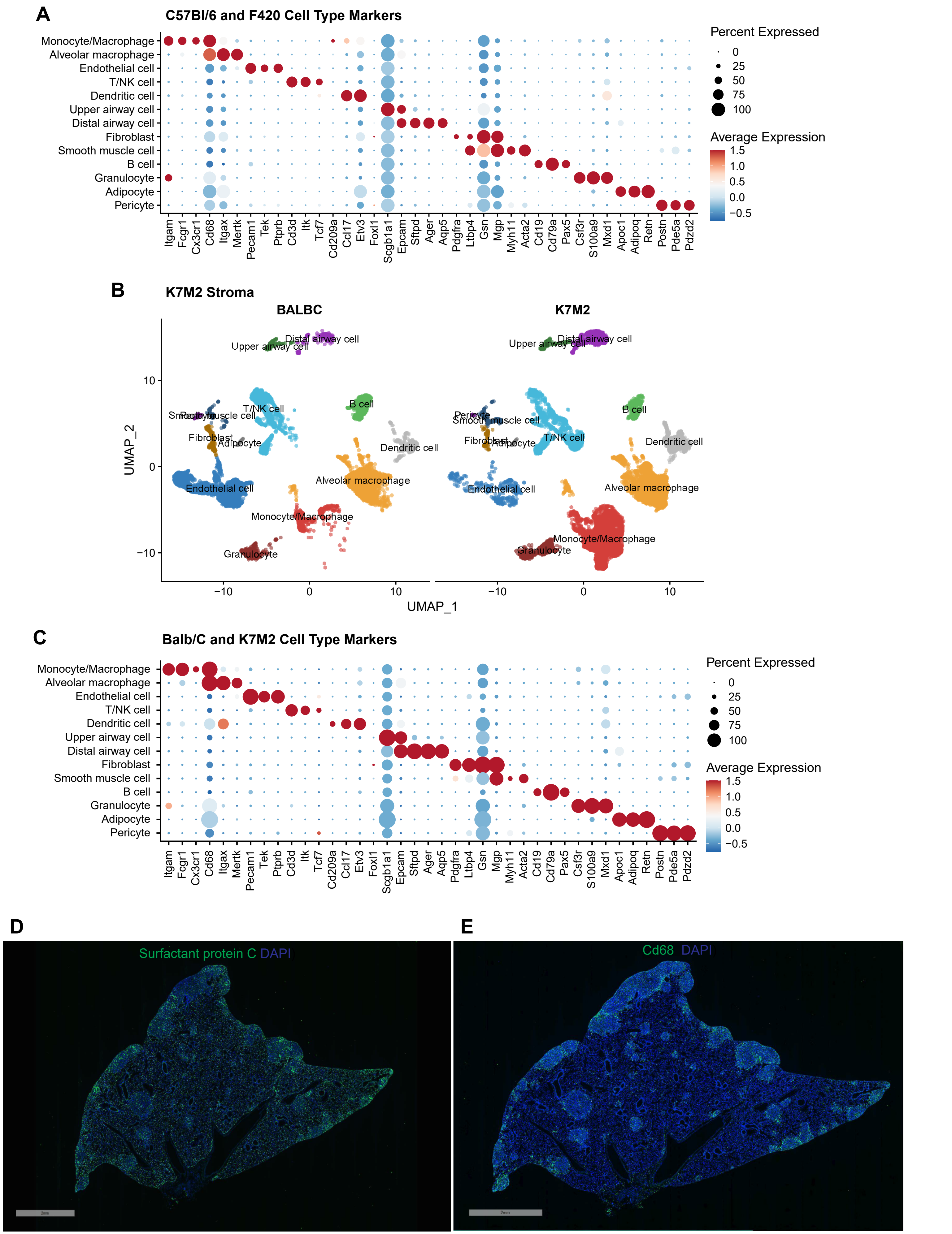

**Supplemental Figure 2:** scRNA-seq marker expression and protein level validation of macrophage and epithelial cell enrichment. **A,** Dotplot demonstrating key marker expression for stromal cell clusters in F420 and CB7Bl/6 samples. **B-C,** UMAP, and dotplot of K7M2 and Balb/C demonstrating stromal clusters and marker expression, respectively. **D-E,** Whole slide scanning IHC confirming metastasis enrichment of surfactant protein C expressing distal airway cells and Cd68 expressing macrophages, respectively. Nuclei counterstained with DAPI in blue. Scale bar= 2mm.

**Supplemental Figure 3: Chronic wound epithelial cells (KRT8+) accumulate in human patient osteosarcoma lung metastasis**

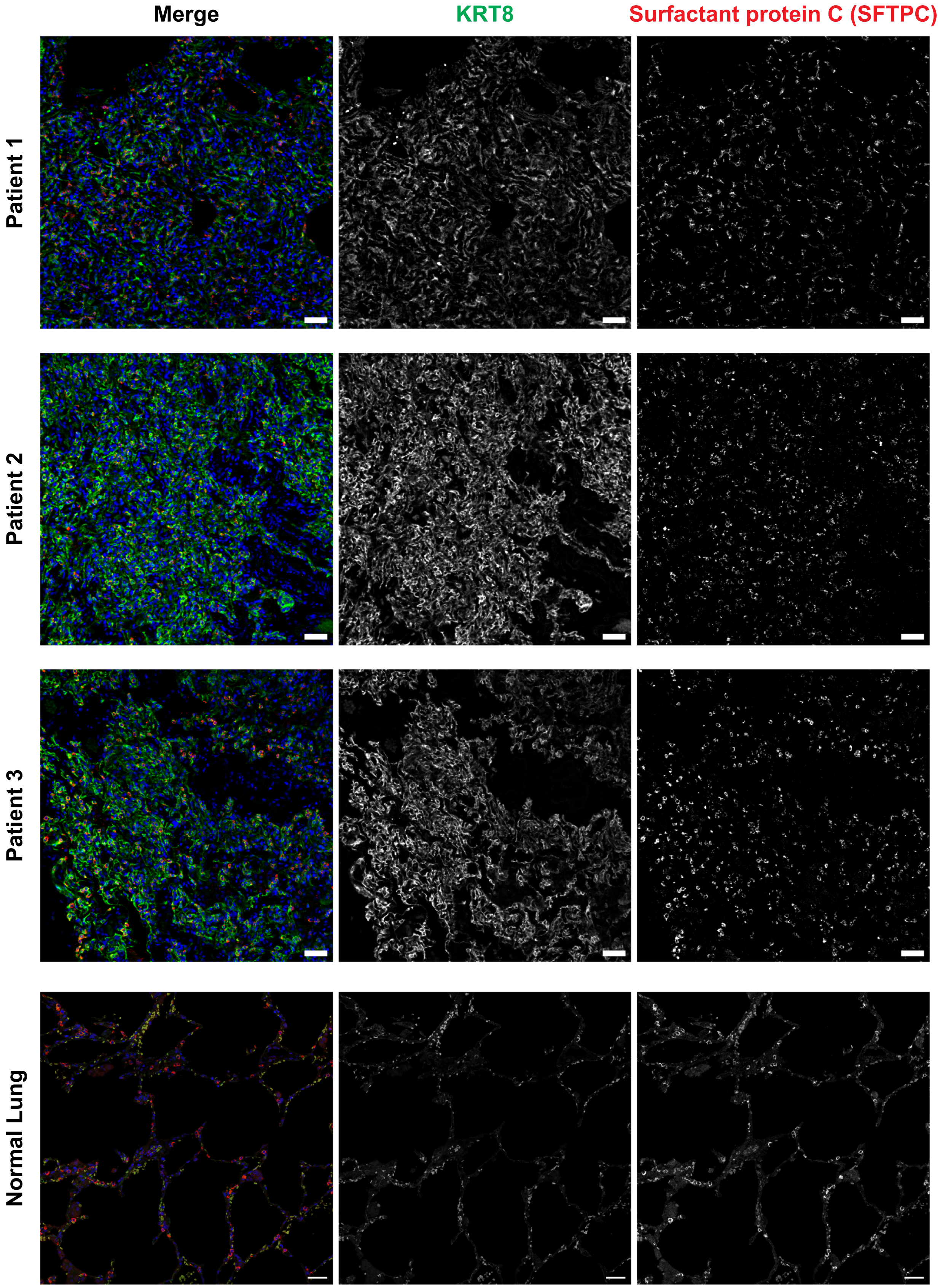

**Supplemental Figure 3:** Chronic wound epithelial cells (KRT8+) accumulate in human patient osteosarcoma lung metastasis. Representative images demonstrating enrichment of KRT8+ DATP (green) and AEC2+ SFTPC in three distinct patient samples compared to uninvolved (normal) lung (from patient 1). Nuclei are counterstained with DAPI in blue. Scale bar= 50µm.

**Supplemental Figure 4: Tumor cell-intrinsic effects of nintedanib**

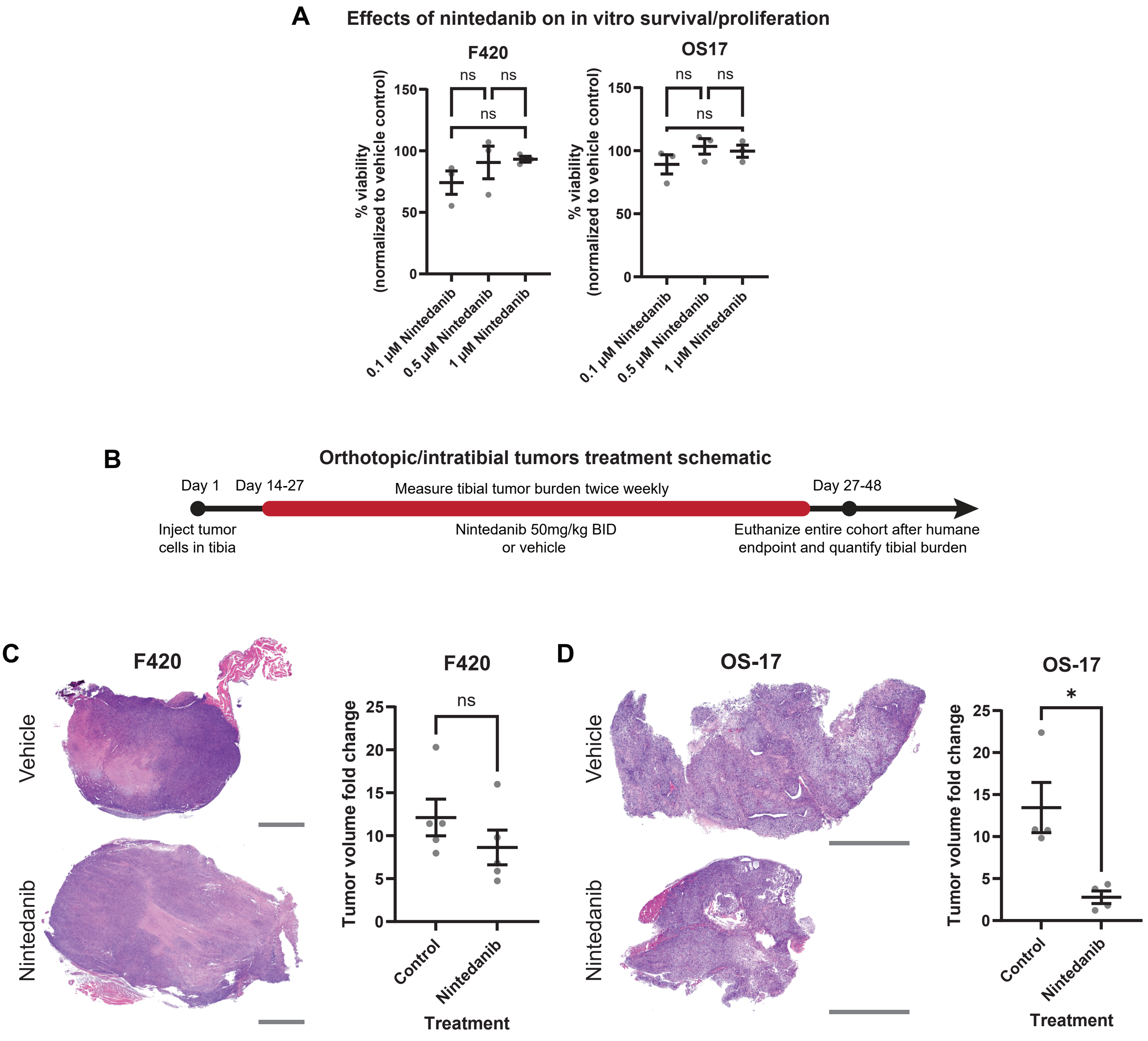

**Supplemental Figure 4:** Tumor cell-intrinsic effect of nintedanib on cell viability and tibial tumor growth. **A,** F420 and OS17 cells were treated *in vitro* with indicated concentrations of nintedanib for 48h and cell viability determined with AlamarBlue assay. N=3. NS=not significant, Brown-Forsythe and Welch's ANOVA with Games-Howell's multiple comparisons test. **B,** Treatment schematic for treatment of tibial tumors with nintedanib *in vivo.* **C-D,** Representative images and quantitation of change in tibial tumor volume in vehicle control compared to nintedanib treated mice in F420 and OS17 models. N=5 mice/treatment group. *$p<0.05$, Welch's T-test. Scale Bar= 1mm.

### Supplemental Figure 5: Ligand-Receptor-Target (NicheNET) gene expression analysis

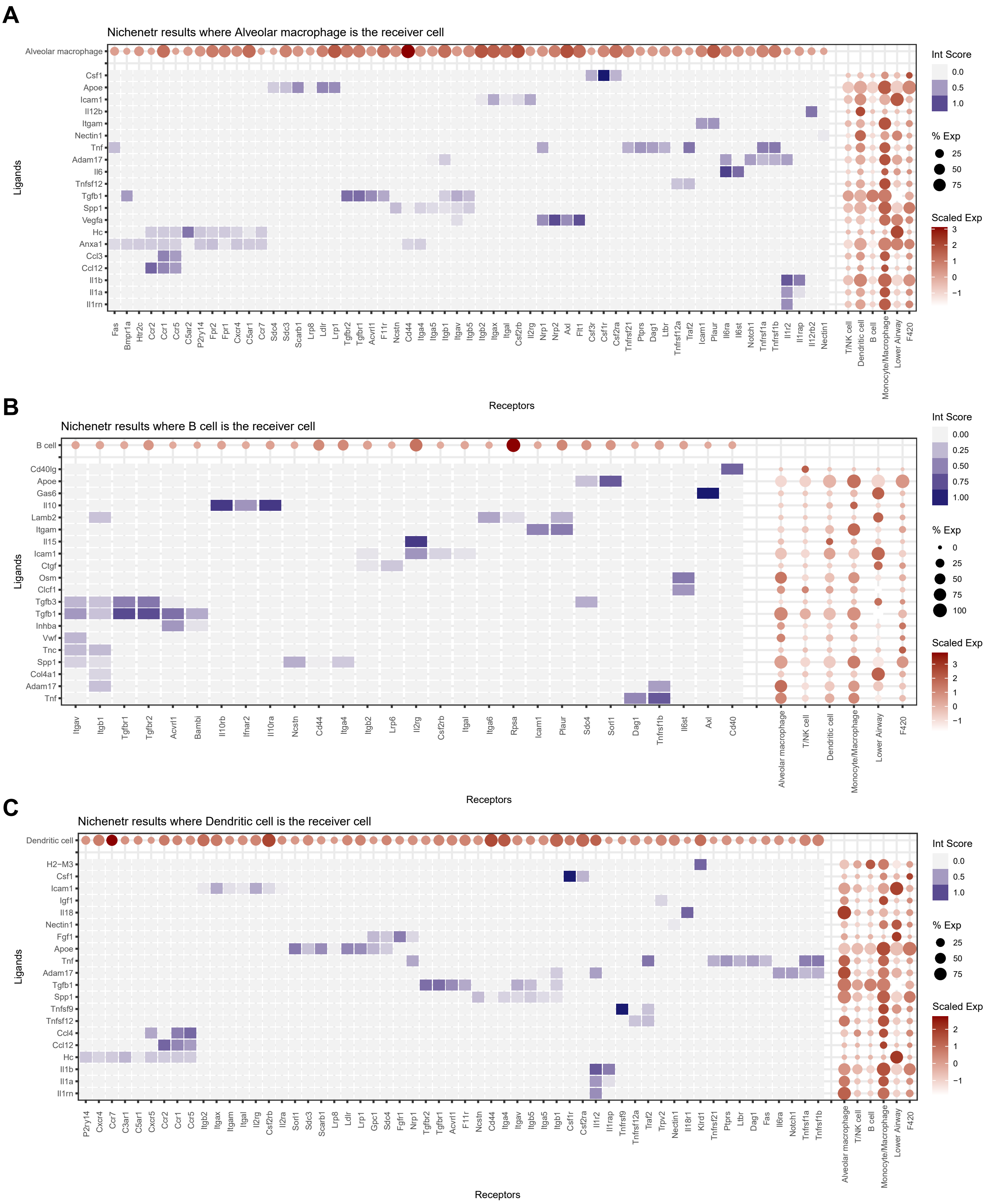

**Supplemental Figure 5: Ligand-Receptor-Target (NicheNET) gene expression analysis**

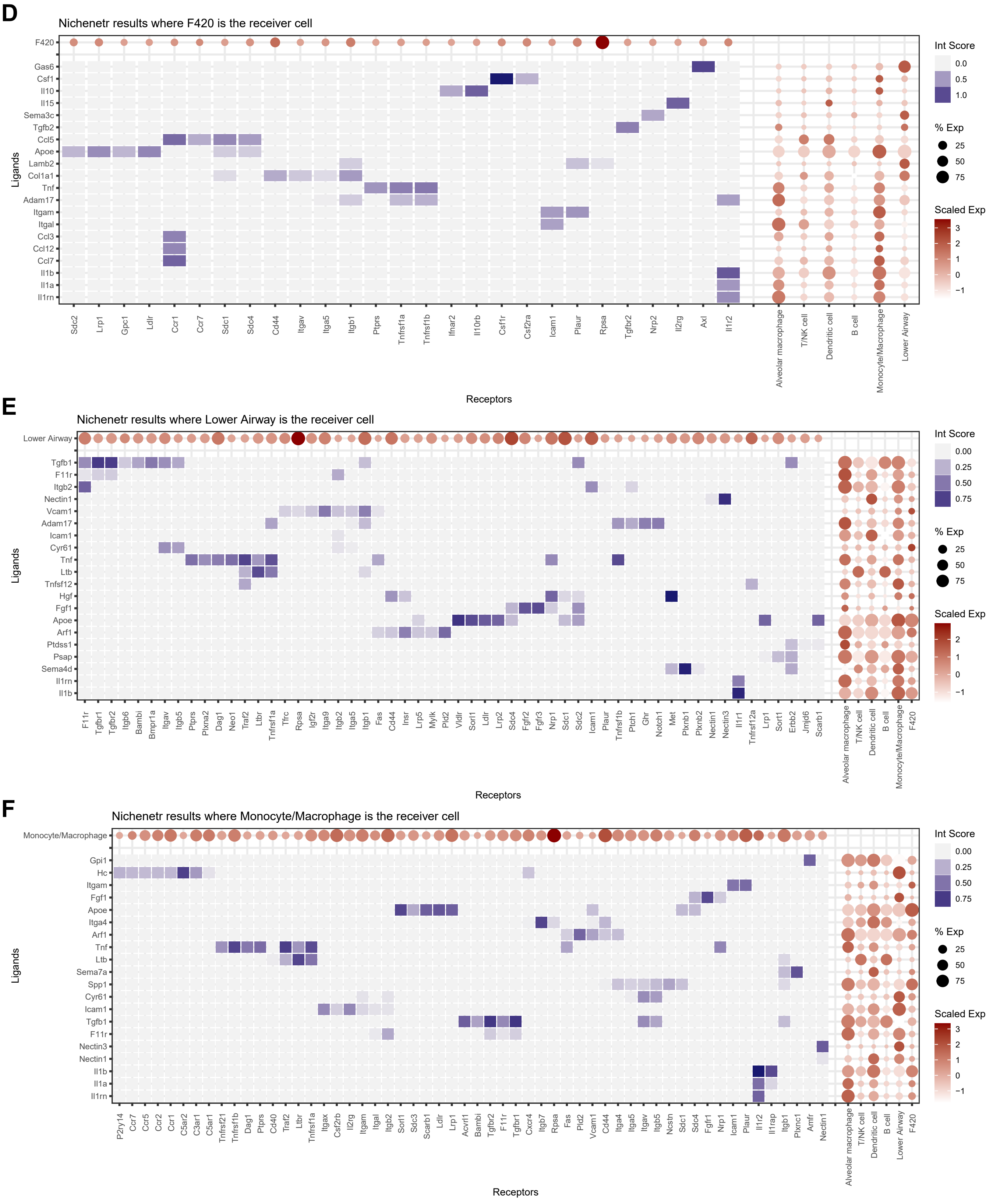

### Supplemental Figure 5: Ligand-Receptor-Target (NicheNET) gene expression analysis

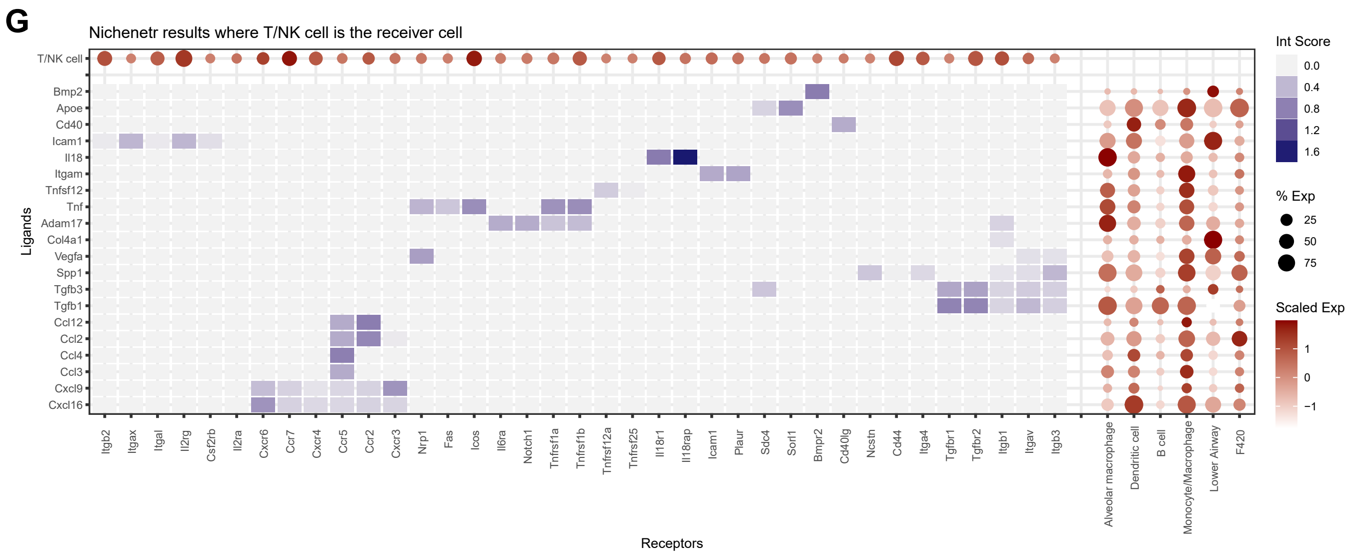

**Supplemental Figure 5:** Ligand-Receptor-Target (NicheNet) cell-cell interactions within metastatic niche. **A-G,** Combined heat map and dotplots showing NicheNet ligand-receptor expression analysis between cells within the metastatic niche.

**Supplemental Figure 6: Immediate vs delayed treatment, tumor burden-based randomization**

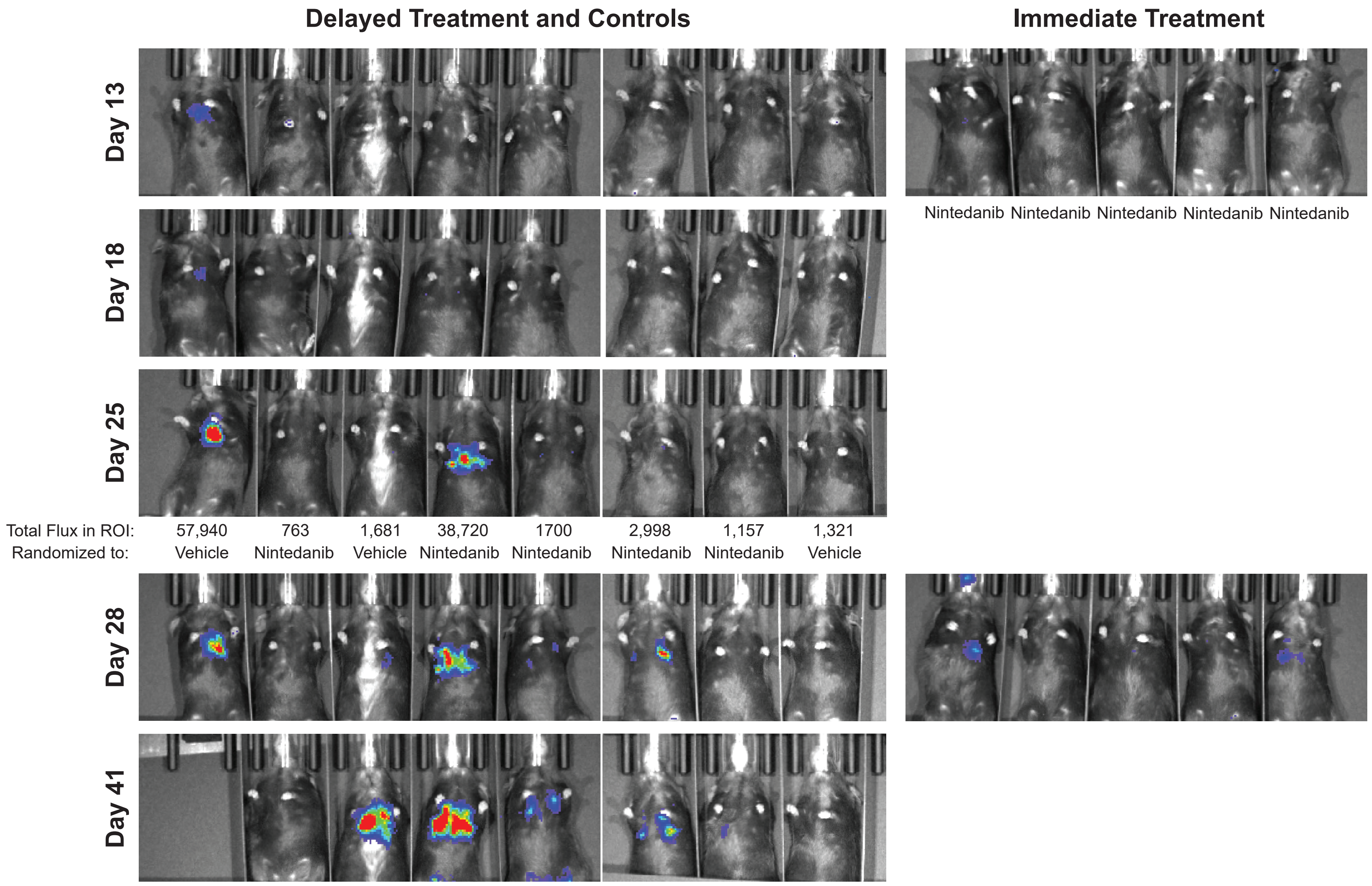

**Supplemental Figure 6:** *In vivo* bioluminescence imaging demonstrating metastasis burden over time in mice treated with vehicle or nintedanib immediately after lung seeding (24 hours post tail-vein injection) or delayed (28 days post tail-vein injection). Relates to Figure 6C.

**Supplemental Figure 7: Mechanistic and pharmacodynamic effects of nintedanib**

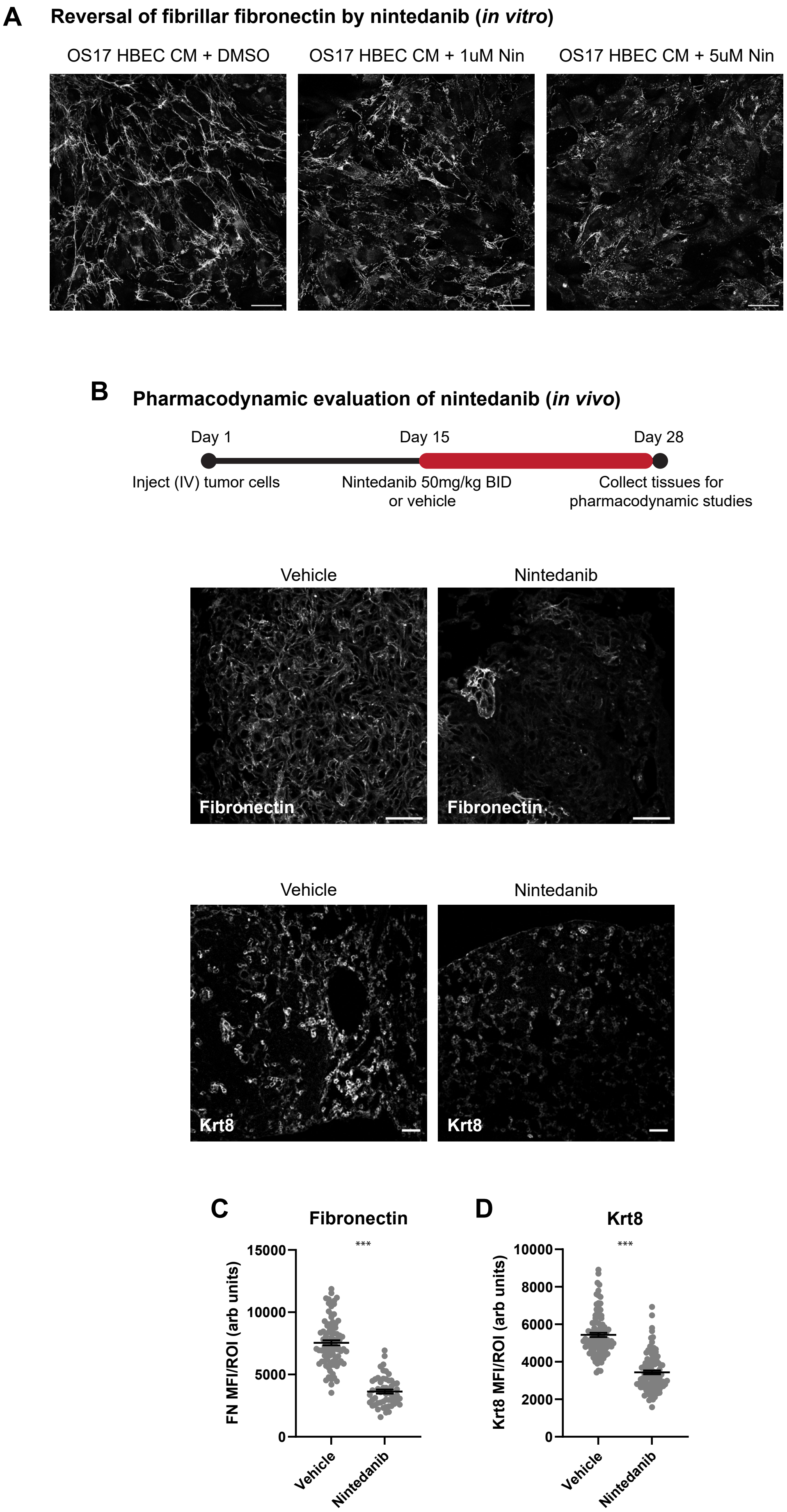

**Supplemental Figure 7:** Protein level validation of key transcriptomic data. **A,** Nintedanib inhibits epithelial-induced fibronectin deposition in OS17 cells. Scale bar= 50μm. **B,** Treatment schematic for nintedanib *in vivo* study on established metastasis for **C,D,** representative IHC images and quantitation for fibronectin and Krt8 in vehicle and nintedanib treated mice. n=3 animals, at least 10 regions of interest per animal. Welch's t-test $p<0.0001$.

**Supplemental Figure 8: Niche-wide changes in target signaling pathway activity with immediate and delayed nintedanib**

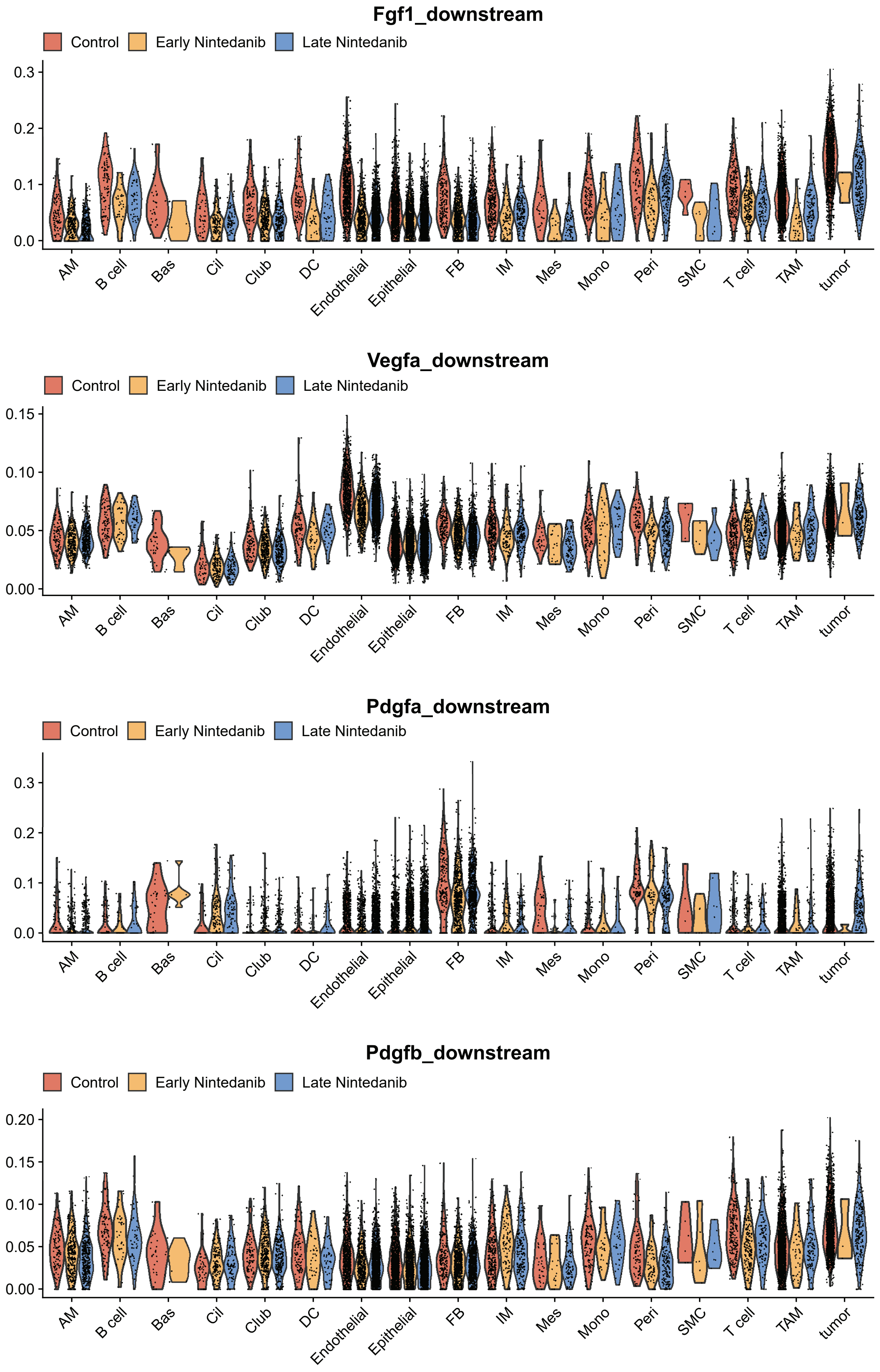

**Supplemental Figure 8:** Niche-wide changes in target signaling pathway activity with immediate and delayed nintedanib treatment. Quantitation by AUC of downstream activation of nintedanib target pathways (Fgf1, Vegfa, Pdgfa, and Pdgfb) in all niche cells. Corresponds to clusters in stromal UMAP in Figure 6D. Note expected downregulation of Vegfa signaling in nintedanib treated endothelial cells while pathway is not impacted in other niche cells.
